# Quantitative metabolic reference for healthy human cerebrum derived from group averaged 9.4T ^1^H MRSI data

**DOI:** 10.1101/2024.04.04.587927

**Authors:** Andrew Martin Wright, Theresia Ziegs, Anke Henning

## Abstract

Drawing inspiration from previous works using ^1^H FID MRSI, this study quantifies metabolite concentrations at 9.4 T in the human cerebrum of a volunteer cohort and performs a respective group analysis to derive region specific metabolite concentrations. Voxel-specific corrections were performed for both water and individual metabolites, as well as used tissue specific T_1_-relaxation times. Anatomical and magnetic resonance spectroscopic imaging data were collected using MP2RAGE and FID MRSI sequences, and subsequent data underwent a series of preprocessing techniques. Results showed consistent metabolite maps for key metabolites (NAA, tCr, Glu, tCho and mI), while instability in data quality was noted for lower slices. This study not only showcases the potential of metabolite quantification and mapping at 9.4 T but also underscores the necessity for meticulous data processing to ensure accurate metabolite representations. Comparisons with earlier works and single voxel results validate the methodologies adopted.

**Highlights:**

- Quantitative whole cerebrum metabolite maps in the human brain acquired at 9.4 T.
- Group averaged regional metabolite concentrations.
- Group summed regional metabolite spectra.

## 1. Introduction

Molecular imaging of the human brain using proton magnetic resonance spectroscopic imaging (^1^H MRSI) offers a unique opportunity to explore metabolite distributions across the human brain as well as estimate concentrations of these metabolites in multiple brain regions. The development of ^1^H MRSI capabilities is a continual process of technical development marked by reducing acquisition times(Bogner et al., 2020) while also improving image resolutions (Nassirpour et al., 2016), enhancing spectral resolutions, and providing reproducible and accurate results (Gasparovic et al., 2006; Hangel et al., 2021; Maudsley et al., 2009; Wright, Murali Manohar, et al., 2021; Ziegs et al., 2023). Improvements in ^1^H MRSI have emphasized its value as a diagnostic tool for neurodegenerative diseases such as multiple sclerosis (Heckova et al., 2022), glioma (Gruber et al., 2017; Hangel et al., 2020; Laino et al., 2020), epilepsy (Pan et al., 2013), and more (Maudsley et al., 2020; Öz et al., 2014).

In order to facilitate a wider-spread use of ^1^H MRSI in clinical decision making, establishing guidelines (i.e. a respective metabolic reference atlas) for typical metabolite concentration ranges and their respective spatial distribution in the healthy brain versus different brain disorders is required. Numerous single voxel ^1^H MRS investigations in the healthy human brain and different brain disorders have been performed but are typically limited to few pre-selected brain regions and impacted by mixing grey and white matter as well as functionally distinct cortices into the same spectroscopy voxel. Previous pioneering work by (Maudsley et al., 2009) derived whole brain metabolite concentration distributions for N-acetyl-aspartate (NAA), choline containing compounds (Cho) and Creatine (Cr) from a group analysis of high-resolution whole brain ^1^H MRSI data acquired at 3 T in 80 healthy volunteers.

Previous studies using ^1^H MRSI at ultra-high field (UHF, field strength ≥ 7 T) strength have demonstrated the potential to map a larger number of brain metabolites at high spatial resolution exploiting the advantages of free induction decay (FID) ^1^H MRSI sequence that minimizes SAR and reduces acquisition time for fully sampled data (Bogner et al., 2020). ^1^H FID MRSI at UHF has unambiguous advantages including increased spectral resolution and increased SNR, or these advantages can be traded for faster acquisition times (Bogner et al., 2012; Henning, 2017; Henning et al., 2009; Nassirpour, Chang, & Henning, 2018; Nassirpour, Chang, Avdievitch, et al., 2018). The ^1^H FID MRSI approach has been successfully translated to a 9.4 T human whole-body MRI scanner and enabled reliable data quality (Nassirpour et al., 2016). Furthermore, 9.4 T studies have investigated full brain reproducibility (Ziegs et al., 2023) and single slice quantitative analysis (Wright et al., 2022) of ^1^H MRSI of the human brain at 9.4 T.

This work melds previous 9.4 T ^1^H FID MRSI efforts to provide group averaged metabolite distribution maps across the human cerebrum as metabolic reference. A respective dedicated data analysis pipeline has been developed and serves as a trail blazer toward the establishment of a metabolic atlas based on larger data sets in future. The same multi-slice 2D acquisition technique was used in this work as an extension of prior 9.4 T ^1^H FID MRSI studies (Nassirpour et al., 2016; Wright et al., 2022; Ziegs et al., 2023); a simulated relaxation corrected macromolecule baseline was included in metabolite fitting (Wright, Murali-Manohar, et al., 2021); and full T_1_-corrections were utilized to correct for T_1_-weighting that accompanied the short TR of this work (Wright et al., 2022). We combined MRSI data sets and respective individual metabolite maps from different volunteers into group averaged ones for the first time at 9.4 T to showcase how MRSI could be utilized in clinical studies where comparisons between healthy controls and diseased cohorts are researched. We hence demonstrate the potential of 9.4 T MRSI in elucidating the quantitated neurochemical profiles in the human cerebrum and provide reference metabolite concentrations for different brain regions.

## 2. Methods

10 healthy volunteers were measured on a 9.4 T whole body scanner (Siemens Magnetom, Erlangen, Germany). A local ethics committee approved acquisition of data in human volunteers, and all volunteers provided written consent prior to scanning. Volunteers were asked to participate for as long as comfortable with a maximum allowable scan duration of two hours inside of the scanner. 8 data sets were useable for analysis; both excluded data sets were due to subject motion which yielded incomplete and non-analyzable data sets. All volunteers were measured using the same dual-row (18Tx/32Rx) phased array head coil (Avdievich et al., 2018).

### 2.1 Acquisition

High-resolution (0.6 x 0.6 x 0.6 mm^3^) anatomical MP2RAGE images were acquired at the beginning of each scan session (TI1/TI2 = 900/3500 ms, FA=4/6°,TR = 6 ms, TE = 2.3 ms, TA = 11 min). Full details of the sequence are described by (Hagberg et al., 2017) with a matched fast AFI map (Pohmann & Scheffler, 2013; Yarnykh, 2007) (resolution: 3.3 x 3.3 x 3.3 mm^3^, FA = 50°, TE/TR_1_/TR_2_ = 4/20/100 ms, TA = 1 min 20 sec) to correct for B_1_ ^+^ inhomogeneity related distortions.

Metabolite and water reference data were acquired with a ^1^H FID MRSI sequence using an acquisition delay (TE*) of 1.3 ms, a TR of 300 ms, a flip angle of 47°, and elliptical k-space shuttering. Metabolite spectra utilized an optimized water suppression scheme with three unmodulated Hanning-filtered Gaussian pulses (BW = 180 Hz, duration = 5 ms) with flip angles of 90°, 79.5°, and 159°; the inter-pulse delay between all pulses was 20 ms. MRSI and water reference data were acquired with a 6 x 6 x 6 mm^3^ resolution. For a slice dimension of 210 x 210 x 6 mm^3^ the acquisition time (TA) was approximately 4.5 min; resulting in approximately 9 min per slice. 2^nd^-order vendor-implemented image based B_0_ shimming was used in blocks to reduce the overall scan time. In most cases, four blocks were sufficient to B_0_ shim the full brain: three blocks with 3 to 5 slices each dividing the cerebrum and a fourth block with 4 to 6 slices for cerebellum data.

Full brain coverage including the cerebellum was planned during data acquisition. However, due to insufficient data quality in lower slices, this study is limited to the metabolic analysis of the cerebrum only. Poor data quality in lower brain slices has also been reported by an earlier study (Ziegs et al., 2023) while using the same RF coil setup and B_0_ shimming method at 9.4T and can be attributed to insufficient SNR for metabolite data due to a limitation of the longitudinal RF coil coverage along with broad linewidth due to insufficient B_0_ shimming. The total slices acquired for each volunteer ranged from 11 to 17 slices depending on the head size and shape.

### 2.2 Preprocessing

^1^H MRSI data were reconstructed using a spatial Hanning filter before the 2D FFT, then eddy current corrected (Klose, 1990), and coil combined using the singular value decomposition (SVD) method (Bydder et al., 2008). Data were then further processed to remove residual water using the Hankel-Lanczos (HLSVD) (Cabanes et al., 2001) method with 10 decaying sinusoids in the range of 4.4-5 ppm. 1^st^-order phase correction was performed using a linear back prediction of the missing FID points (Kay, 1988). L2-regularization for retrospective skull lipid removal (Bilgic et al., 2014) was performed uniformly on all data sets using findings from (Ziegs et al., 2023). Spectroscopic imaging data from individual volunteers was then fitted as outlined in section 2.3.

### 2.3 Group averaged metabolite maps

Data were fitted in LCModel (Provencher, 2001) from 0.8 to 4.2 ppm using a simulated basis set with 12 metabolites : N-acetyl-aspartate (NAA), total creatine (tCr), aspartate (Asp), *γ*-aminobutyric acid (GABA), taurine (Tau), glutamine (Gln), glutamate (Glu), glutathione (GSH), myo-inositol (mI), scyllo-inositol (Scyllo), N-acetyl-aspartyl-glutamate (NAAG), and phosphocholine + glycerophosphocholine (tCho) plus a simulated MM spectrum (MM_AXIOM_, (Wright, Murali-Manohar, et al., 2021; Wright, Murali Manohar, et al., 2021)) to account for MM contributions. After spectral fitting, data were quantified using a voxel-specific correction method considering relaxation correction of metabolites and water as described in (Wright et al., 2022) yielding metabolite maps in mmol kg^-1^ (Figure 1) and mM (presented in Supporting Information).

**Figure 1:**
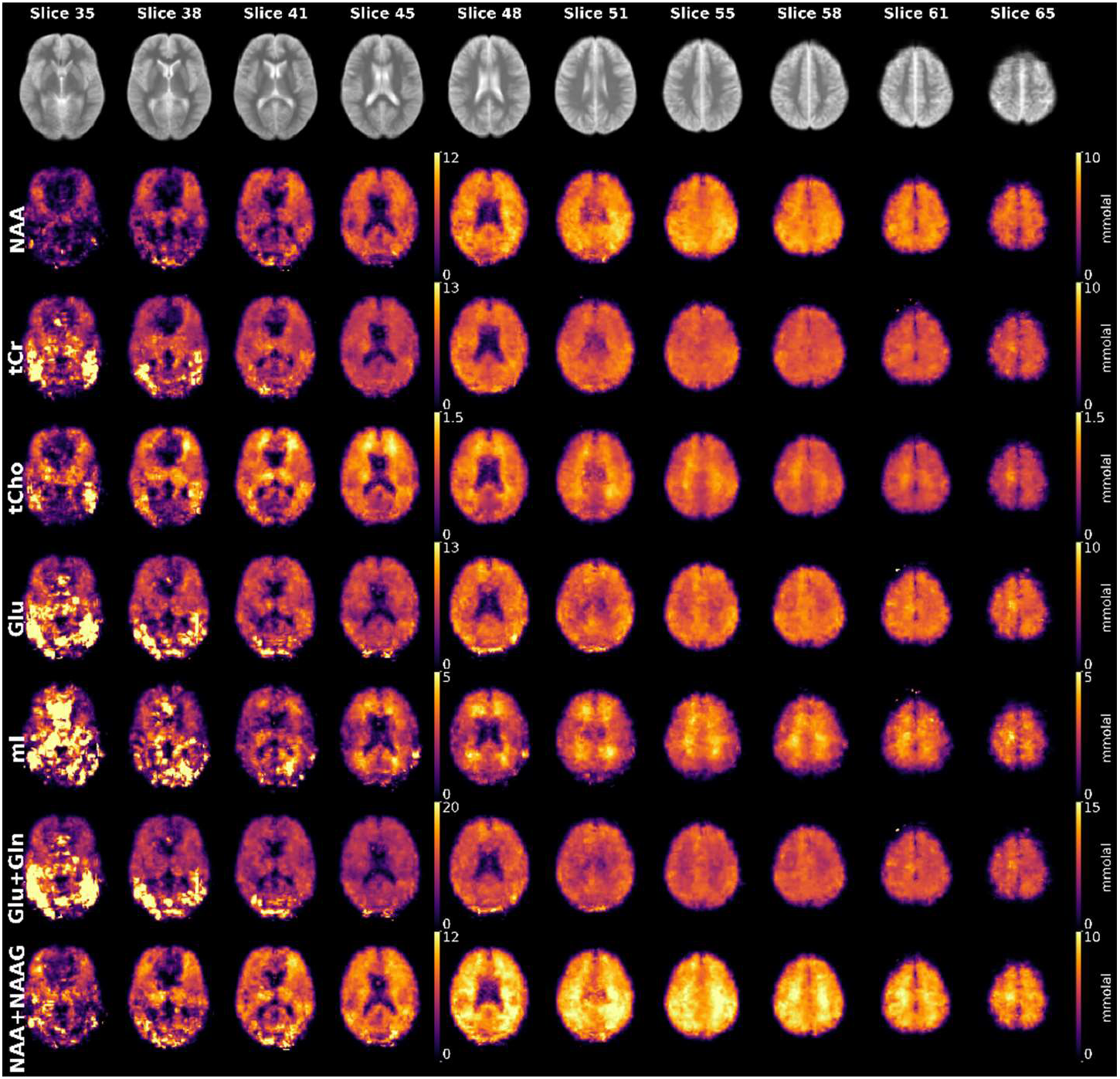
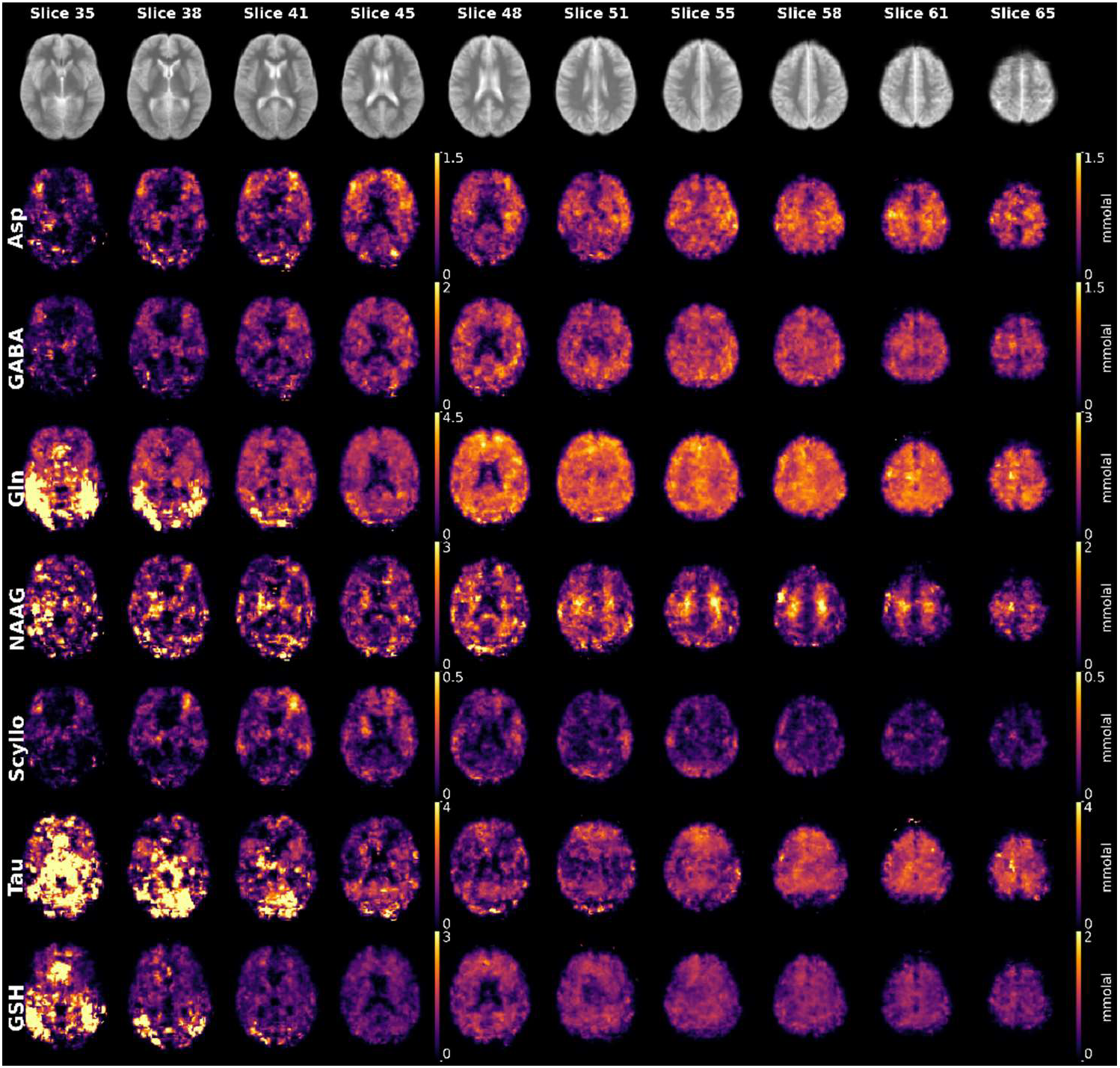
Median metabolite maps of eight volunteers displaying the distribution of all analyzed metabolites. The anatomical data are derived from averaged, T1-weighted images coregistered to the MNI 152 Human Brain Atlas. The quantitative data, expressed in mmol kg^-1^, are also coregistered to the same atlas. Slice positions refer to the slice position in MNI space.

Following quantification, quantitative metabolite maps were interpolated to match the MP2RAGE spatial resolution and written to the nifti file format (.nii) as preparatory step for linear co-registration calculations to be performed in FSL (Jenkinson et al., 2012). These quantitative metabolite maps were then co-registered to the MNI152 Human Brain atlas (2mm resolution) using FSL FLIRT by transforming the MP2RAGE to the standardized space and having all metabolite maps be secondary transformations. The FLIRTed metabolite maps were then used for calculation of average and median metabolite and CRLB maps.

Brain regions were partitioned using three atlases (Harvard-Oxford maximum probability cortical atlas [2 mm], Harvard-Oxford maximum probability subcortical atlas [2 mm], and Johns Hopkins University, International Consortium for Brain Mapping template [2mm], (Desikan et al., 2010; Frazier et al., 2005; Goldstein et al., 2007; Hua et al., 2008; Makris et al., 2006; “MRI Atlas of Human White Matter,” 2006; Wakana et al., 2007)) and used to calculate concentrations for eight regions in the brain. A full summary of regional combinations is reported in Supporting Information Table 1.

**Table 1:**
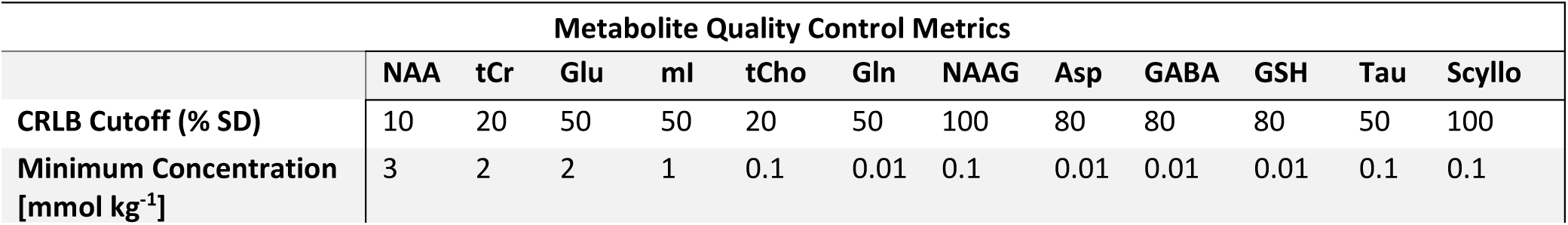
Maximum CRLB cutoffs and minimum concentrations for metabolite concentration estimates. The CRLB thresholds were defined as the mean plus SD for each individual metabolite as recommended by the MRS consensus group (Lin et al., 2021).

### 2.4 Brain region specific average spectra and metabolite concentrations

Eight anatomical regions are reported in this study. These regions were defined by taking atlas data and combining sub-regions into the following major brain regions: frontal lobe GM, frontal lobe WM, parietal lobe GM, parietal lobe WM, occipital lobe GM, occipital lobe WM, temporal lobe GM, and corpus callosum. These combinations are explicitly reported in Supporting Information Table 1 and are shown in Supporting Information Figure 1.

Regional summed spectra were created after transforming the regional masks back to the original space for each volunteer. This was done using the FSL suite and using the inverse of the affine matrix from the forward FLIRT calculation. These masks were then resampled back to the original MRSI resolution (6 x 6 x 6 mm^3^), and the indices for non-zero voxels were then used to select spectral data. The .coord output files were used to create the summed spectra. Summed spectra were then evaluated across all slices, and slices that contained little signal or strong lipid distortions were excluded from the summed spectra.

Mean regional concentrations were calculated using quantified metabolite maps following the FSL FLIRT operation. To reduce the impact of spurious spectral fitting, data were filtered for maximum allowable CRLB values and minimum physiological concentrations as a quality measure (Table 1). Concentrations were calculated in mmol kg^-1^ and mM quantities. Concentrations between regions were compared for each metabolite. Mann-Whitney U-tests were carried out for all reported regions with a significance threshold of 0.05 assigned. Bonferroni corrections were applied for each region and used to report corrected p-values for comparisons.

## 3. Results

### 3.1 Quantitative Metabolite Maps

Metabolite maps with T_1_-corrected and quantitated data for 12 metabolites are reported in Figure 1. The calculated maps are the median from eight volunteers and reported in MNI space with units of mmol kg^-1^. Metabolite map slices (in MNI space) from 45 and below are reported with an additional scale to reduce oversaturation for some metabolite maps. Metabolite maps in mM quantities are reported in Supporting Information Figure 2.

**Figure 2:**
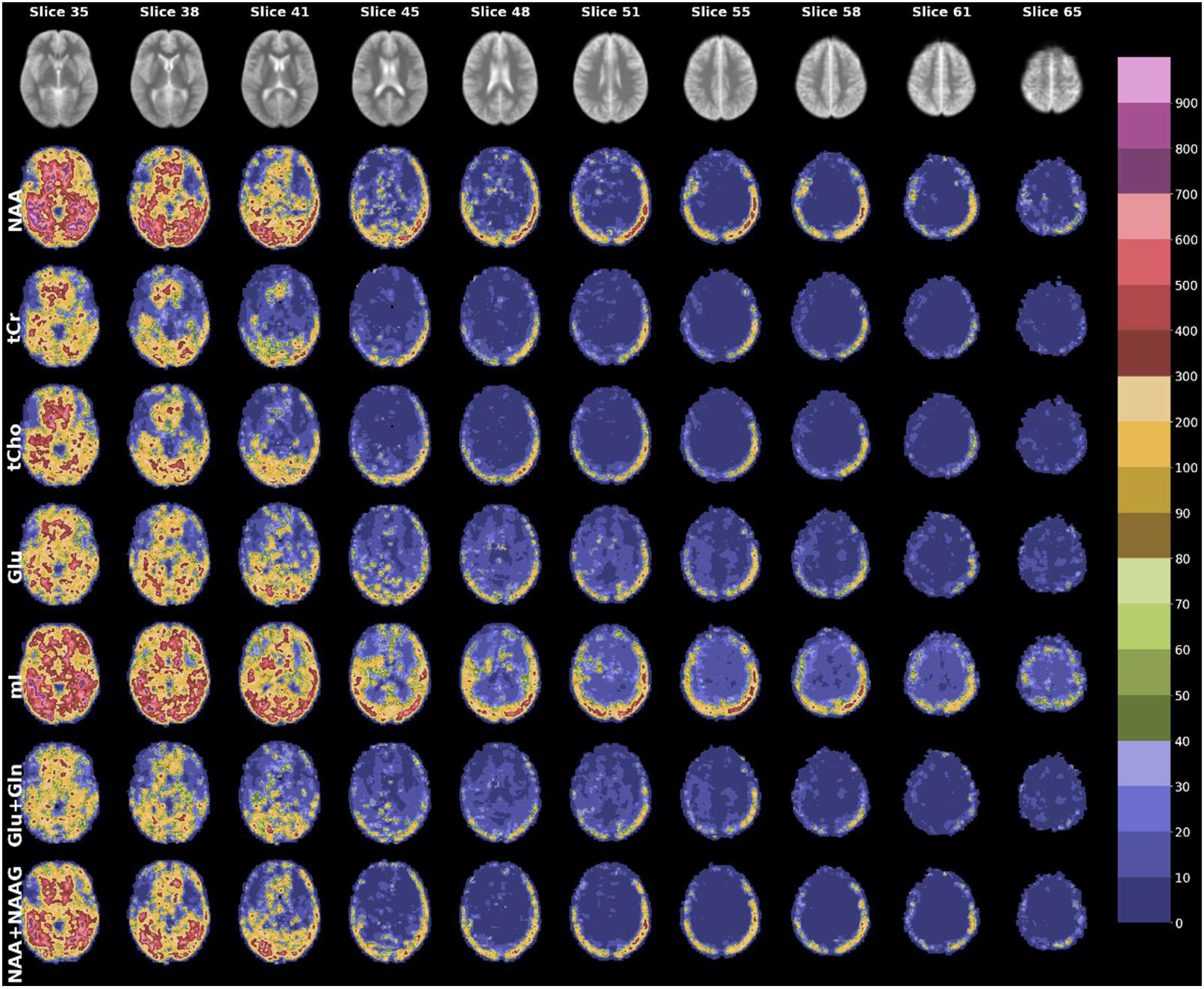
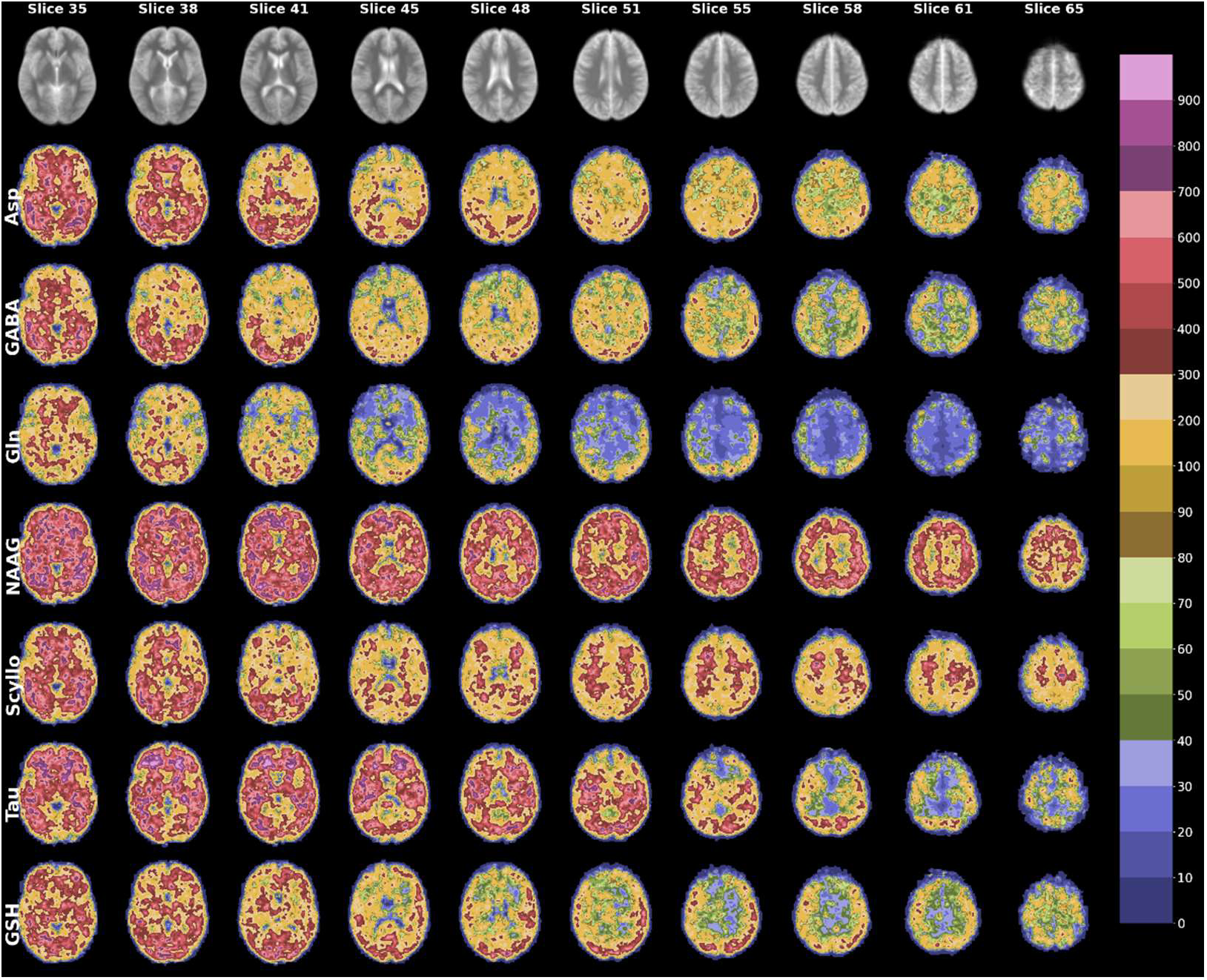
median CRLB (% SD) maps with a non-uniformly binned color bar. These maps do not include the threshold which was applied for metabolite concentration calculations. Maps with this threshold applied are shown in Supporting Information Figure 3. All maps are presented in the MNI 152 space.

Observable tissue contrast is apparent for tCho, Glu, Gln, mI, NAAG, Glu+Gln, and NAA+NAAG maps. The distribution of metabolite concentrations is consistent with previous 9.4 T reports (Nassirpour et al., 2016; Wright et al., 2022; Ziegs et al., 2023) and lower field strengths (Hangel et al., 2021; Maudsley et al., 2009).

Averaged CRLB maps are shown in Figure 2. The color bar is a non-uniformly spaced binning of CRLB ranges. CRLB maps with maximum thresholds applied are reported in Supporting Information Figure 3. As can be seen in Figure 1 and Figure 2, data in MNI space below slice 45 diminish in quality. CRLB maps show that the median CRLB in lower slices is much higher than acceptable for data reporting. This effect is seen clearly in Figure 1 by areas of signal dropout or overexposure. Metabolite maps presented in sagittal, coronal, and transversal are also reported (Figure 6) and showcased in section 4.1.

**Figure 3:**
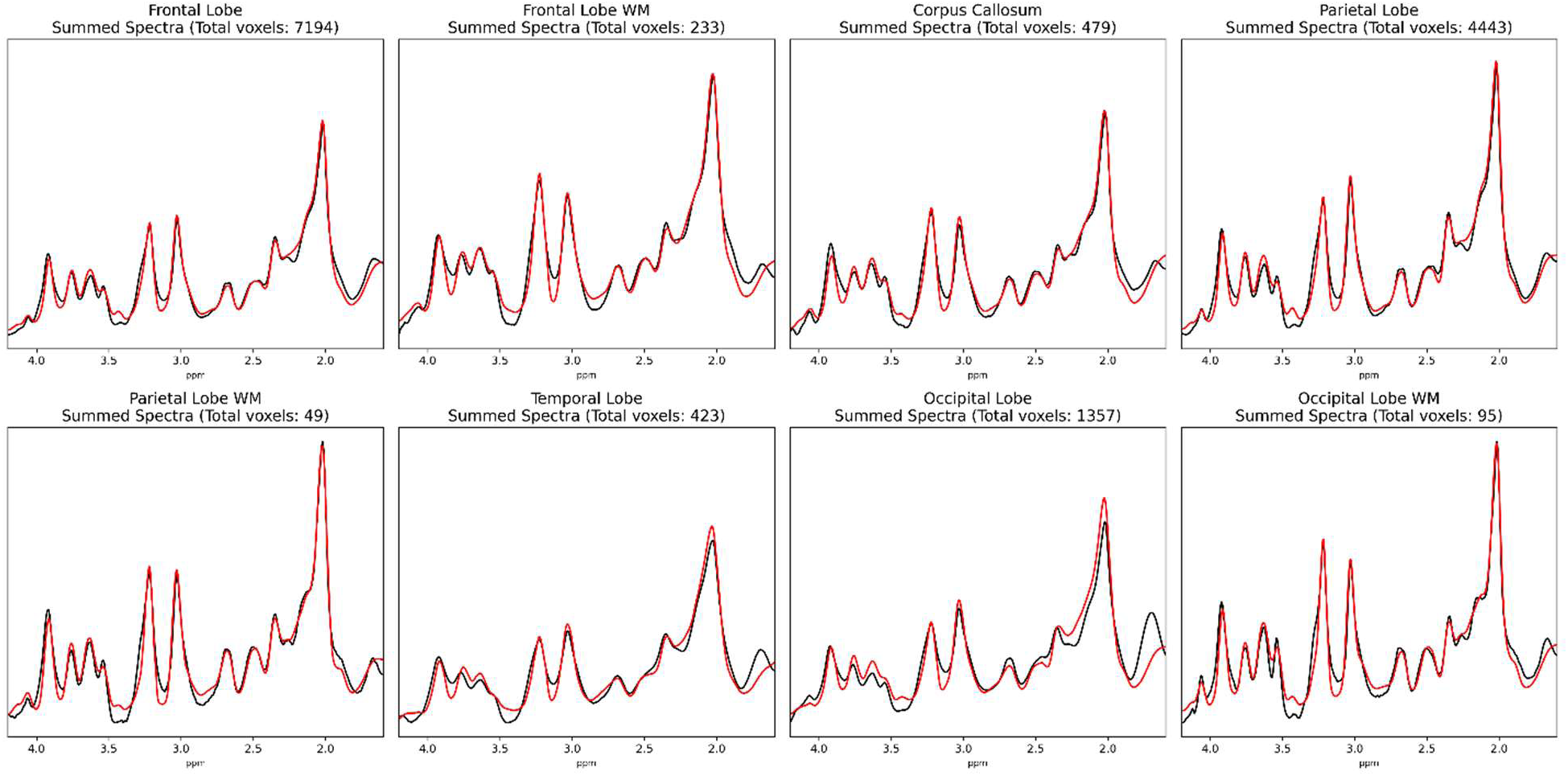
Summed metabolite spectra for each of the eight chosen major brain regions. Data were summed by taking the .coord output files. The black line represents the actual data, and the red line is the sum of fit results for selected voxels.

### 3.2 Regional Metabolite Spectra

The LCModel fits of metabolites were combined to display regional sums for metabolite spectra (Figure 3) where the black line is the spectra data and the red line is the fit data. Estimated regional spectra and concentrations include the following anatomical regions: frontal Lobe GM, frontal Lobe WM, parietal Lobe, parietal Lobe WM, temporal lobe, insular cortex, corpus callosum. All regional masks applied to MNI space images were transformed back to each original space and MRSI resolution and all non-zero signal voxels were summed. The total number of voxels included for each region is listed below the region name in Figure 3.

All regions maintain good spectral resolution. However, regions with more voxels (specifically GM regions) have slightly broader lineshapes.

### 3.3 Regional Metabolite Concentrations

Metabolite concentrations [mmol kg^-1^] for eight brain regions are reported in Table 2 and statistical differences marked by numerical superscripts to note significance between regions (Regional Key for statistical comparisons, Table 3) and visually represented in Figure 4. Metabolite concentrations were calculated in the MNI152 space by masking quantitative metabolite maps with a regional mask. Concentrations are in general agreement with previous work at 9.4 T (Nassirpour et al., 2016; Wright et al., 2022; Ziegs et al., 2023). However, there are some slight differences that are discussed further in section 4.2 with violin plots (Figure 5) to show distributions of metabolite concentrations taken from the masked quantified metabolite maps.

**Figure 4:**
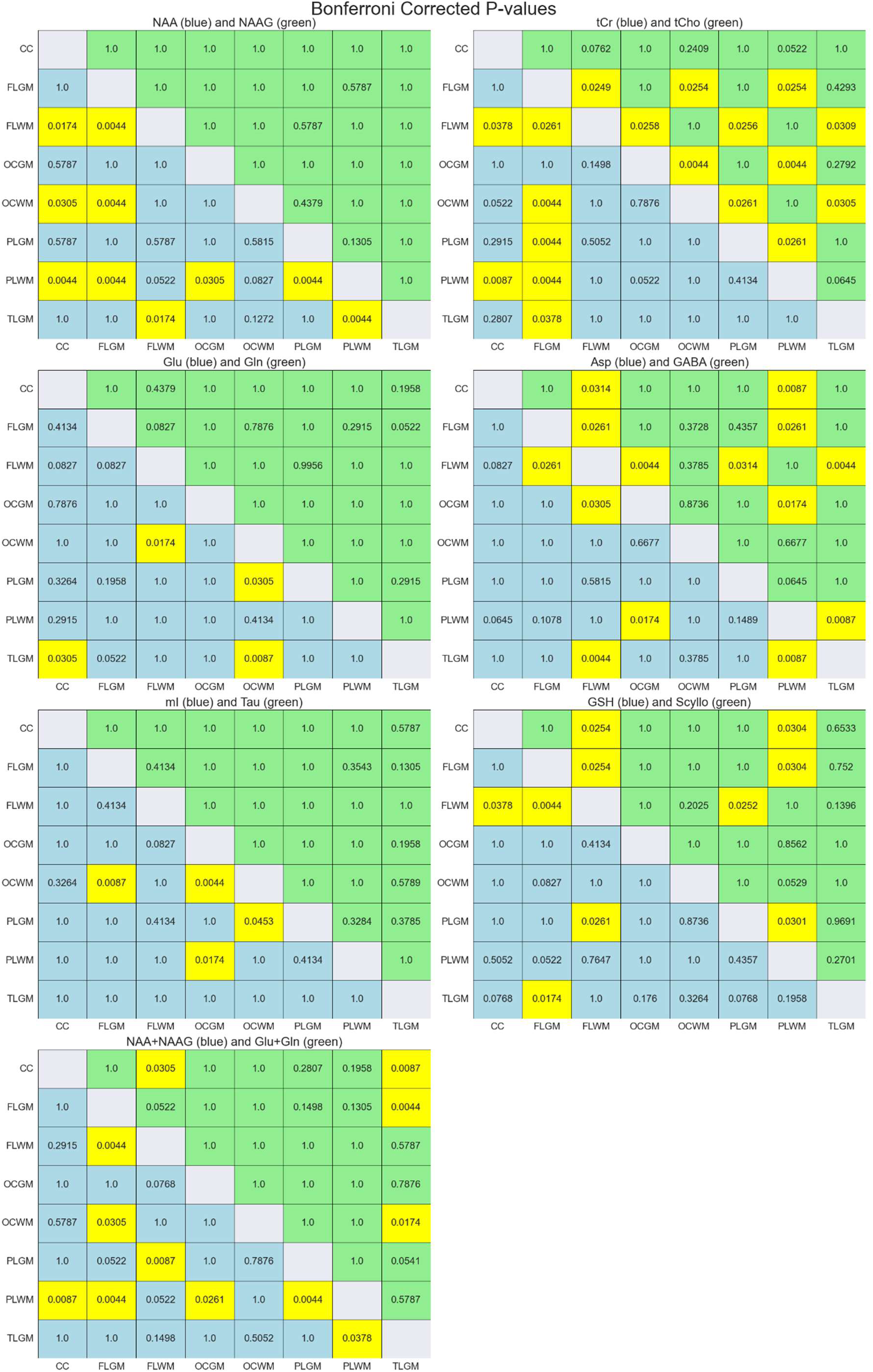
Bonferroni-corrected p-values for all regions and metabolites analyzed. Each heat map reports two metabolites, one below the diagonal (blue) and one above the diagonal (green). Corrected p-values below 0.05 are represented with yellow shaded cells. **CC**: corpus callosum, **FLGM**: frontal lobe grey matter, **FLWM**: frontal lobe white matter, **OCGM**: occipital lobe grey matter, **OCWM**: occipital lobe white matter, **PLGM**: parietal lobe grey matter, **PLWM**: parietal lobe white matter, **TLGM**: temporal lobe grey matter

**Figure 5:**
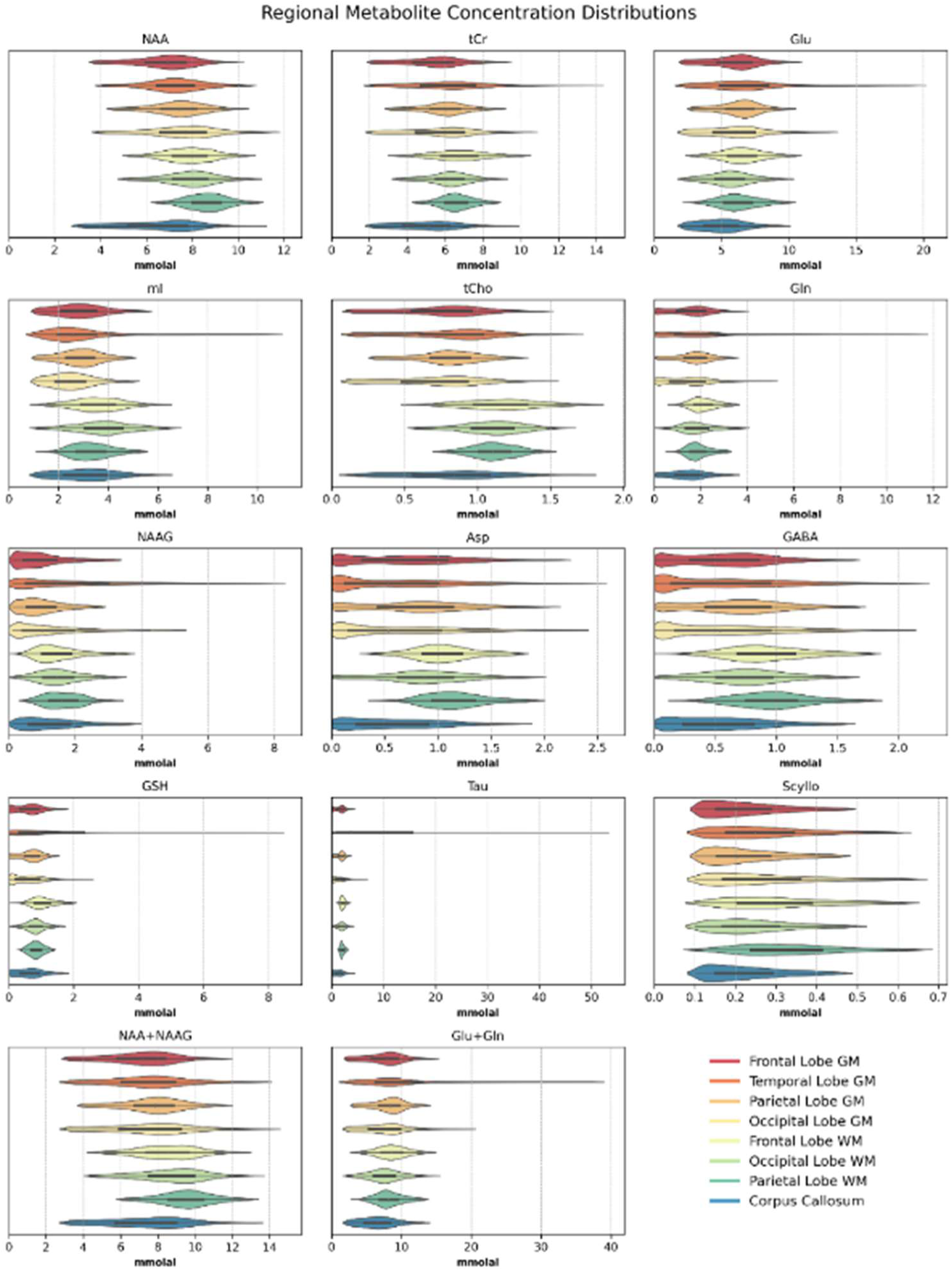
violin plots showing regional metabolite concentrations for eight major brain regions. Data had a CRLB threshold applied (Table 1). These distributions were aggregated in MNI space by applying the regional masks across all volunteers.

**Table 2:** Metabolite concentrations [mmol kg^-1^] for the selected eight major brain regions. Regional masks can be found in Supporting Information Figure 1. Metabolite concentrations reported in mM quantities are reported in Supporting Information Table 2. The superscript on metabolites highlights an adjusted p-value < 0.05 between regions as is defined in the Regional Key for Statistical Comparisons (Table 3).

| Metabolite Concentrations [mmol kg<sup>-1</sup>] |  |  |  |  |  |  |  |  |
| --- | --- | --- | --- | --- | --- | --- | --- | --- |
|  | Frontal Lobe GM | Temporal Lobe GM | Parietal Lobe GM | Occipital Lobe GM | Frontal Lobe WM | Occipital Lobe WM | Parietal Lobe WM | Corpus Callosum |
| **NAA**<sup>2,3,7,8,16,17,19,21,24,28</sup> | 9.39 ± 0.73 | 9.99 ± 0.66 | 10.23 ± 0.69 | 10.40 ± 1.20 | 11.16 ± 0.34 | 11.27 ± 0.69 | 12.4 ± 0.62 | 8.64 ± 1.47 |
| **tCr**<sup>2,3,8,12,17,23</sup> | 7.57 ± 0.32 | 10.76 ± 6.33 | 8.48 ± 0.37 | 8.02 ± 0.74 | 9.54 ± 0.66 | 8.91 ± 0.68 | 9.50 ± 0.79 | 7.15 ± 1.08 |
| **Glu**<sup>5,15,22,26</sup> | 8.41 ± 0.39 | 14.15 ± 12.98 | 9.07 ± 0.41 | 8.94 ± 1.48 | 9.50 ± 0.76 | 8.05 ± 0.40 | 9.34 ± 1.51 | 7.24 ± 1.27 |
| **ml**<sup>8,10,19</sup> | 4.06 ± 0.49 | 8.65 ± 8.89 | 4.14 ± 0.41 | 3.62 ± 0.46 | 5.24 ± 0.96 | 5.63 ± 0.99 | 5.03 ± 0.77 | 4.36 ± 0.71 |
| **tCho**<sup>3,6,8,10,13,15,17,19,21,24,26</sup> | 1.08 ± 0.08 | 1.22 ± 0.12 | 1.14 ± 0.08 | 1.00 ± 0.14 | 1.66 ± 0.11 | 1.57 ± 0.16 | 1.60 ± 0.12 | 1.18 ± 0.22 |
| **Gln** | 2.37 ± 0.26 | 8.01 ± 13.19 | 2.63 ± 0.33 | 2.74 ± 1.08 | 3.22 ± 0.57 | 3.06 ± 0.72 | 3.22 ± 0.97 | 2.29 ± 0.59 |
| **NAAG** | 1.78 ± 0.92 | 2.80 ± 1.30 | 1.65 ± 0.54 | 2.07 ± 0.79 | 3.09 ± 1.49 | 2.47 ± 0.60 | 2.92 ± 1.05 | 2.26 ± 0.96 |
| **Asp**<sup>3,6,19,24,28</sup> | 1.02 ± 0.18 | 0.82 ± 0.26 | 1.12 ± 0.27 | 0.80 ± 0.40 | 1.63 ± 0.31 | 1.33 ± 0.38 | 1.71 ± 0.35 | 0.92 ± 0.35 |
| **GABA**<sup>2,3,6,13,16,17,19,24,28</sup> | 0.86 ± 0.08 | 0.85 ± 0.16 | 1.00 ± 0.10 | 0.83 ± 0.21 | 1.38 ± 0.16 | 1.11 ± 0.27 | 1.44 ± 0.27 | 0.83 ± 0.20 |
| **GSH**<sup>2,3,13,23</sup> | 0.96 ± 0.12 | 3.51 ± 3.84 | 1.03 ± 0.16 | 1.08 ± 0.33 | 1.63 ± 0.34 | 1.3 ± 0.20 | 1.28 ± 0.20 | 1.00 ± 0.23 |
| **Tau** | 2.64 ± 0.51 | 22.16 ± 26.51 | 2.76 ± 0.32 | 2.78 ± 0.91 | 5.94 ± 7.16 | 3.41 ± 1.38 | 4.25 ± 2.01 | 7.50 ± 14.68 |
| **Scyllo**<sup>2,3,13,16,17,21</sup> | 0.30 ± 0.04 | 0.36 ± 0.04 | 0.31 ± 0.03 | 0.37 ± 0.10 | 0.44 ± 0.05 | 0.29 ± 0.07 | 0.49 ± 0.09 | 0.28 ± 0.06 |

**Table 3:** Key to statistical comparisons noted in Table 2 and visually displayed in Figure 4. For example, the number 1 represents a comparison between the Frontal Lobe GM and Corpus Callosum and the number 19 represents a comparison between the Parietal Lobe WM and Occipital Lobe GM.

| Regional Key for Statistical Comparisons |  |  |  |  |  |  |  |  |
| --- | --- | --- | --- | --- | --- | --- | --- | --- |
| Region | Key Numbers |  |  |  |  |  |  |  |
| Corpus Callosum |  |  |  |  |  |  |  |  |
| Frontal Lobe GM | 1 |  |  |  |  |  |  |  |
| Frontal Lobe WM | 2 | 3 |  |  |  |  |  |  |
| Occipital Lobe GM | 4 | 5 | 6 |  |  |  |  |  |
| Occipital Lobe WM | 7 | 8 | 9 | 10 |  |  |  |  |
| Parietal Lobe GM | 11 | 12 | 13 | 14 | 15 |  |  |  |
| Parietal Lobe WM | 16 | 17 | 18 | 19 | 20 | 21 |  |  |
| Temporal Lobe GM | 22 | 23 | 24 | 25 | 26 | 27 | 28 |  |
| Region | Corpus Callosum | Frontal Lobe GM | Frontal Lobe WM | Occipital Lobe GM | Occipital Lobe WM | Parietal Lobe GM | Parietal Lobe WM | Temporal Lobe GM |

Inter-subject coefficients of variation (CVs) were calculated using mmol kg^-1^ metabolite maps and are reported in Table 4, and corrected p-values from Mann-Whitney U-tests are reported and shown in Figure 4. All p-values less than 0.05 are highlighted in yellow. Regional metabolite concentration distributions are shown in Figure 6 via violin plots and discussed in section 4.2. Please refer to Supporting Information Table 4 for an additional representation of regional metabolite concentration differences.

**Table 4:** inter-subject coefficients of variation (CV, %)

| Inter-Subject CV (%) |  |  |  |  |  |  |  |  |
| --- | --- | --- | --- | --- | --- | --- | --- | --- |
|  | Frontal Lobe GM | Temporal Lobe GM | Parietal Lobe GM | Occipital Lobe GM | Frontal Lobe WM | Occipital Lobe WM | Parietal Lobe WM | Corpus Callosum |
| NAA | 5 | 5 | 4 | 6 | 2 | 5 | 4 | 14 |
| tCr | 3 | 3 | 3 | 8 | 5 | 6 | 5 | 12 |
| Glu | 3 | 3 | 3 | 6 | 5 | 4 | 7 | 11 |
| ml | 9 | 10 | 8 | 12 | 15 | 15 | 10 | 15 |
| tCho | 5 | 6 | 5 | 12 | 6 | 7 | 6 | 16 |
| Gln | 8 | 10 | 8 | 14 | 12 | 17 | 14 | 21 |
| NAAG | 15 | 15 | 11 | 18 | 22 | 16 | 18 | 30 |
| Asp | 13 | 21 | 20 | 30 | 4 | 22 | 12 | 28 |
| GABA | 8 | 12 | 9 | 18 | 7 | 16 | 11 | 19 |
| GSH | 10 | 9 | 12 | 13 | 12 | 10 | 12 | 20 |
| Tau | 7 | 15 | 5 | 9 | 14 | 31 | 8 | 18 |
| Scyllo | 5 | 10 | 7 | 16 | 6 | 17 | 12 | 15 |

## 4. Discussion

### 4.1 Metabolite Maps

The metabolite map quality obtained in this study is a step forward aiming at the establishment of a whole brain metabolite concentration reference atlas. In comparison to a previous group analysis study performed at 3T (Maudsley et al., 2009) in a large volunteer cohort demonstrating the spatial distribution of NAA, tCho and tCr a substantially larger number of metabolites could be imaged across the cerebrum herein. Previous whole-brain 7T ^1^H MRSI studies in single volunteers showed the spatial distribution of only five metabolites: NAA, tCr, tCho, Glx and mI (Hangel et al., 2021) and six metabolites without quantification: NAA, tCr, tCho, Glu, mI, and NAAG (Klauser et al., 2021). Compared to previous 9.4 T MRSI studies (Nassirpour, Chang, Avdievitch, et al., 2018; Nassirpour et al., 2016; Ziegs et al., 2023), this work shows a comparable quality across the cerebrum and maintained the standard of quantifying concentrations for 12 metabolites in vivo with voxel-specific T_1_-weighting corrections. However, the quality of metabolite maps was not sufficient to derive quantitative metabolite maps in lower slices in the brain, which is also evidenced in a recent 9.4 T study (Ziegs et al., 2023) which assessed test-retest reproducibility and brain coverage of ^1^H MRSI with an identical technical study design. This limitation has also been demonstrated in previous 7T whole-brain ^1^H MRSI studies (Hangel et al., 2021; Klauser et al., 2021; Motyka et al., 2019). Improving the longitudinal coverage of transmit and receive B_1_ fields by improved RF coil design along with tackling the B_0_ inhomogeneity in the lower brain by dedicated B_0_ shimming software and hardware are likely the most important targets for achieving whole brain ^1^H FID MRSI coverage at UHF in future.

Metabolite concentration distributions obtained herein agree with previous ^1^H MRSI studies (Bogner et al., 2012; Hangel et al., 2021; Henning et al., 2009; Maudsley et al., 2009; Nassirpour et al., 2016; Wright et al., 2022; Ziegs et al., 2023). Metabolites with concentrations appearing higher in GM in metabolite maps include: Glu, Gln, and Tau, while metabolites with increased WM concentrations include: NAA, tCho, mI, and NAAG. In the previous 9.4 T quantitative ^1^H FID MRSI work (Wright et al., 2022), the NAA distribution was not as apparent in the transversal plane; however, in the sagittal and coronal planes (Figure 4), there appears to be an increased concentration of NAA in WM compared to grey matter. These differences observed in the metabolite maps are substantiated with the statistical testing highlighting increased NAA in the parietal lobe WM compared to the GM of the frontal, parietal, and occipital lobes.

By considering Figure 2, it is possible to ascertain further insights into metabolite distributions. Metabolites with CRLBs that appear with a strong tissue contrast provide a hint that regions with an increase in CRLB are less likely to confidently fit a metabolite due to its lower regional concentration. For instance, NAAG is well documented to be almost exclusively located in the WM in MRSI studies (Bogner et al., 2012; Hangel et al., 2021; Henning et al., 2009; Nassirpour et al., 2016; Wright et al., 2022; Ziegs et al., 2023), and this is strongly supported in the present cohort by both the metabolite and CRLB maps. Metabolites with interesting CRLB map contrasts reflecting GM – WM differences can be observed for NAAG, Gln, Tau, and Scyllo. This information could be beneficial in tailoring fitting basis vectors to include the most relevant metabolite information to improve fitting accuracy.

### 4.2 Regional Spectra and Metabolite Concentrations

This work builds on the process of reporting quantitative metabolite concentrations from ^1^H MRSI data by means of voxel-specific T_1_-corrections. A detailed description of these corrections is reported by (Wright et al., 2022), and has been developed further to format the data so that it can be transformed to standard brain spaces. Previous MRSI studies tend to report results in non-quantitative units, with average T_1_-corrections, and with LCModel estimated MMs (Bogner et al., 2012; Gasparovic et al., 2006; Hangel et al., 2021; Henning et al., 2009; Maudsley et al., 2009; Nassirpour et al., 2016).

Regional spectra have good spectral resolution with particularly sharp peaks in WM regions. The broader lineshapes in GM regions are most likely attributable to the voxels on the periphery of the brain which are more susceptible to B0 inhomogeneities. Additionally, GM regions included significantly more voxels dispersed across a larger spatial region, of which many are on the periphery of the brain. The interior WM regions are thus less susceptible to lipid contributions and are comprised of fewer spectra due to the more compact volume of the available atlases. An atlas partitioned into WM regions for use with spectroscopy data sets would allow for the inclusion of more voxels in WM regional spectra.

Future efforts could control spectral quality in an autonomous fashion by considering the linewidths of key peaks, or by considering a spectrum’s match to a control spectrum using a prediction model. However, this likely would not be necessary, and regional spectra quality (specifically SNR) could be expected to improve by collecting more data and improving B_0_ shimming.

Magnetic resonance spectroscopy studies generally report metabolite concentrations in absolute concentrations in either molal (often, mmol kg^-1^) or molar (often, mM) quantities (Near et al., 2020). Regional concentrations estimated in this work are compared to previous single voxel ^1^H MRS and ^1^H MRSI studies in Table 5 [mmol kg^-1^] and Table 6 [mM]. Metabolite concentrations derived herein are in general agreement with previous studies cited in Table 5 and Table 6.

**Table 5:**
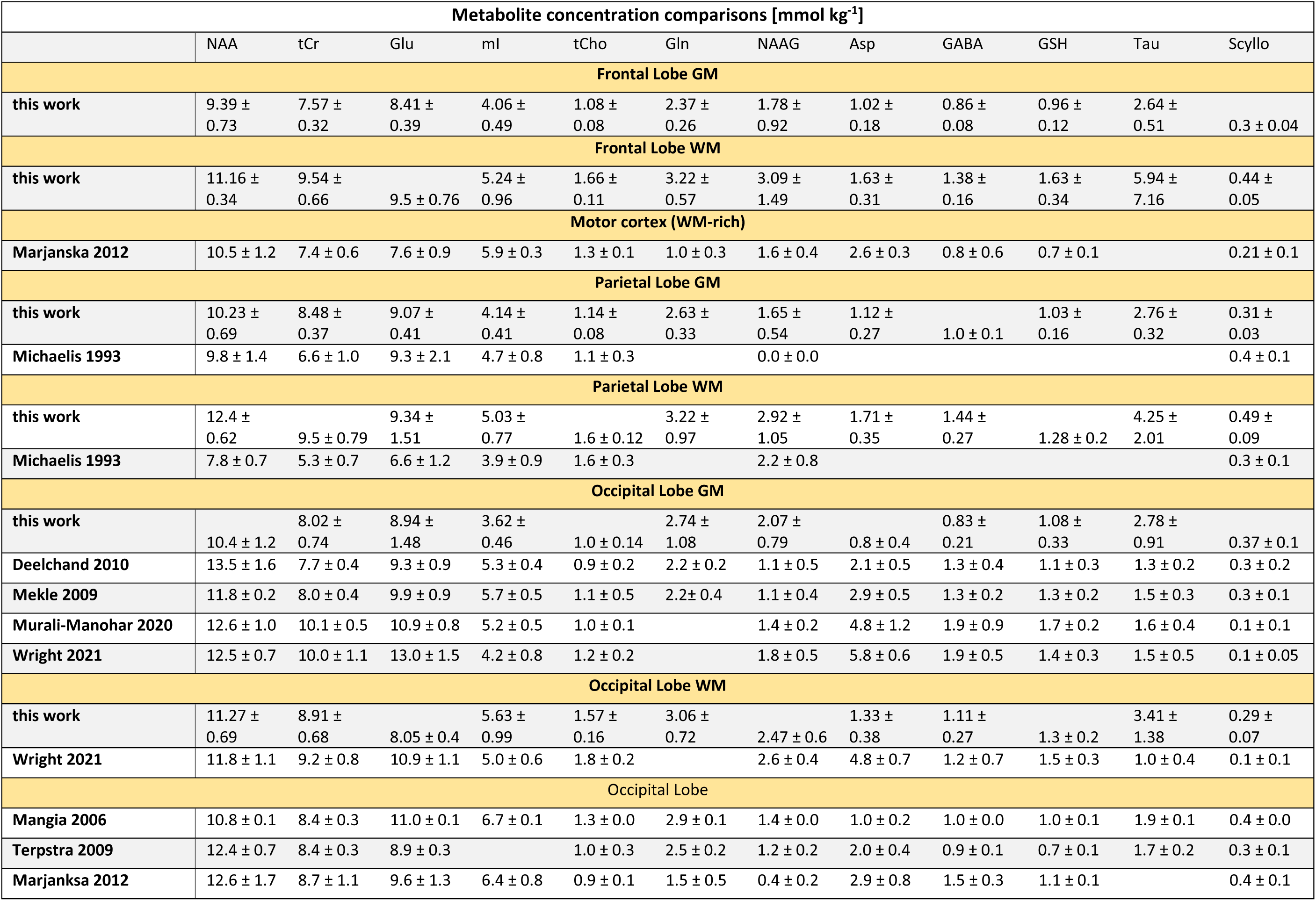
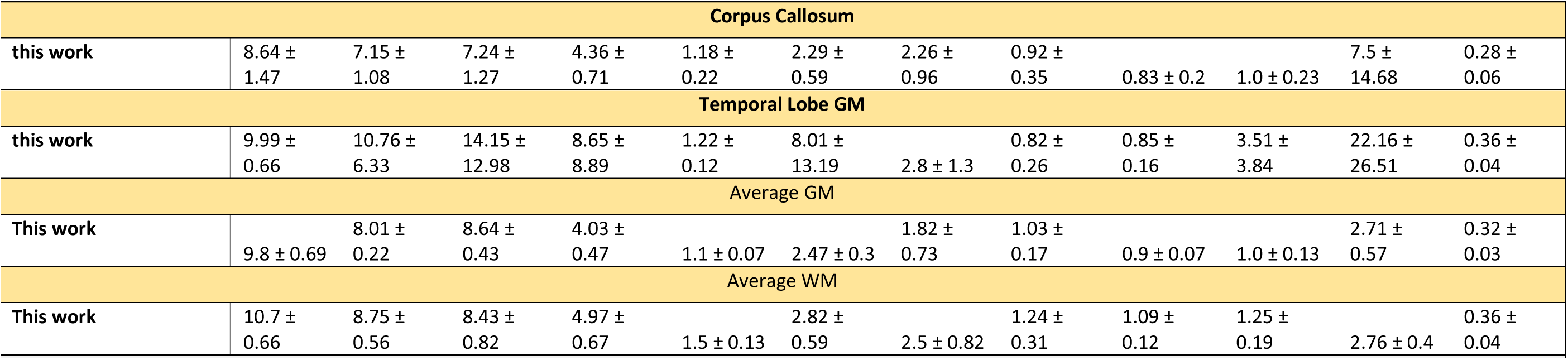
Metabolite concentration comparisons from studies reporting results in [mmol kg^-1^]. Citations: (Deelchand et al., 2010; Mangia et al., 2006; Marjańska et al., 2012; Mekle et al., 2009; Michaelis et al., 1993; Murali-Manohar et al., 2020; Terpstra et al., 2009; Wright et al., 2022; Wright, Murali-Manohar, et al., 2021)

**Table 6:**
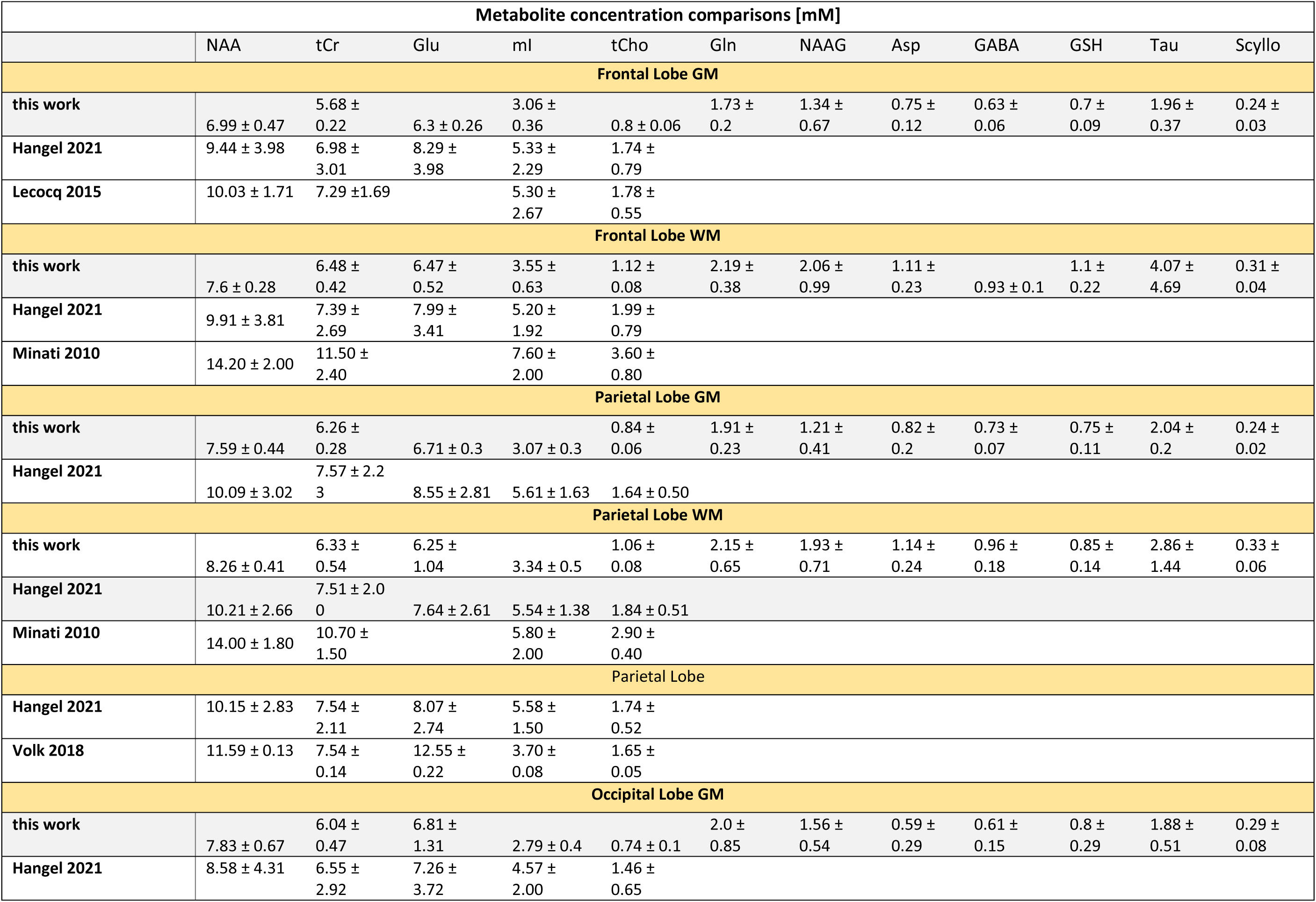
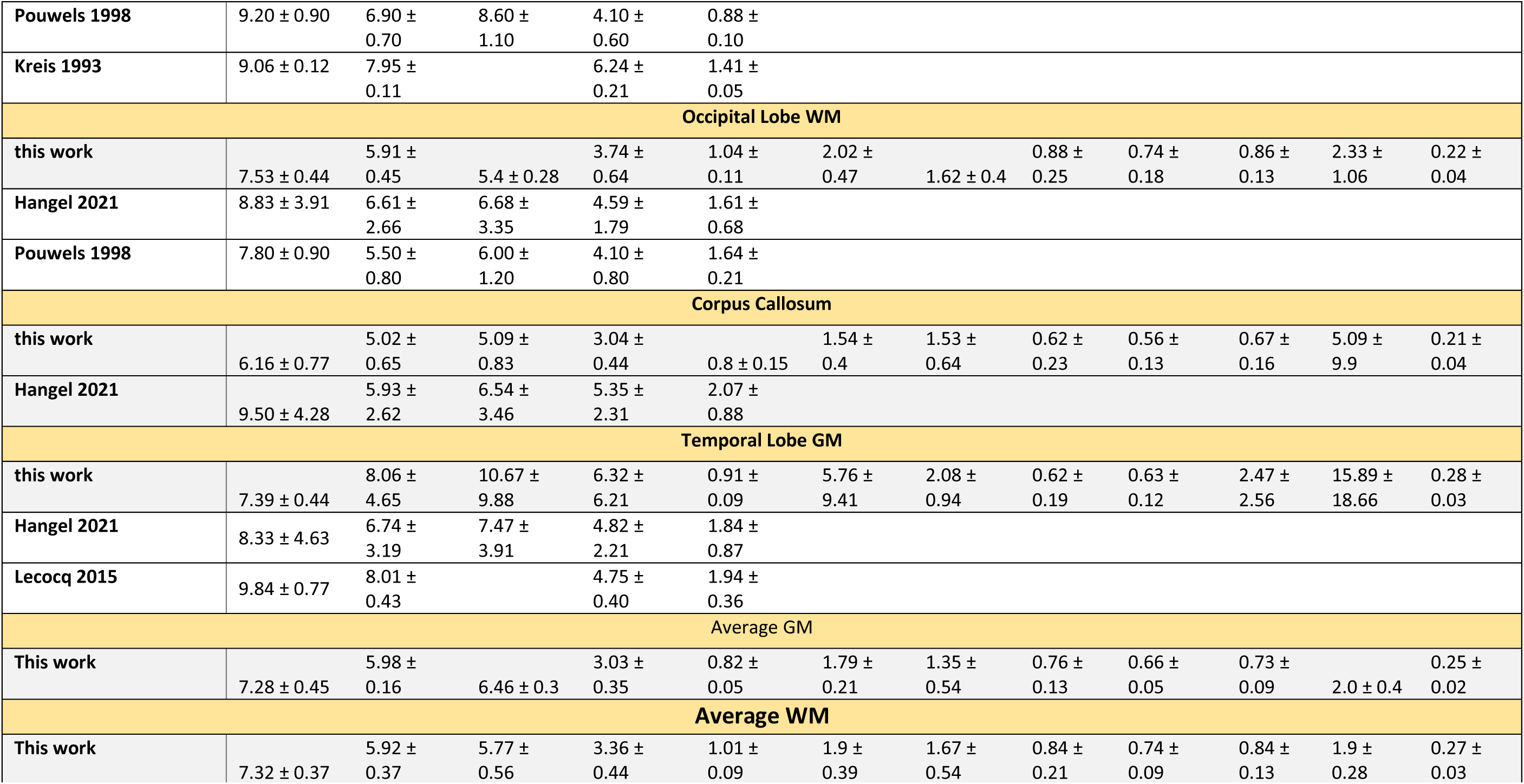
Metabolite concentration comparisons from studies reporting results in [mM]. Citations: (Hangel et al., 2021; Kreis et al., 1993; Le Lecocq et al., n.d.; Minati et al., 2010; Pouwels & Frahm, 1998; Volk et al., 2018)

While metabolite maps show tissue contrast for NAA, tCho, NAAG, Glu, and Gln, metabolite concentrations in Table 2, Figure 4, and Supporting Information Table 2, the differences between GM and WM regions are generally minor if relaxation corrections are considered. NAA, tCho, mI, and Asp are present in higher concentrations in the frontal lobe WM, occipital lobe WM, and parietal lobe WM. In GM regions, it would be expected for Glu and Gln to display and be presented in higher concentrations compared to the concentrations in WM regions. While this is not as obvious for all regions in Table 2, violin plots and metabolite maps (Figure 1) display tissue contrast for Glu and somewhat for Gln; where Glu is elevated in the frontal lobe GM and parietal lobe GM compared to the parietal lobe WM (Table 2 and Figure 4). However, more data and controls for Glu and Gln fitting quality are needed to decisively quantify the concentration differences for Glu and Gln between GM and WM regions.

mI is reduced in GM compared to previous works (Deelchand et al., 2010; Hangel et al., 2021; Kreis et al., 1993; Le Lecocq et al., n.d.; Mekle et al., 2009; Murali-Manohar et al., 2020); this arises from the difference of how these studies accounted for macromolecules. In this work, the MM_AXIOM_ has macromolecule resonances included between 3.4 and 4.0 ppm that result in a reduction of the fitting area for mI, while prior works neglected to include macromolecule resonances which affect the resonances between 3.4 and 4.0 ppm. A similar effect can be seen for NAA, albeit this is not as strong of an effect. Of the available comparisons in Table 5, the metabolite concentrations here tend to be lower. The concentrations are generally overlapping when considering the standard deviations in conjunction with the means between studies. The study from (Minati et al., 2010) reported much higher concentrations than other works. It is also important to note that the herein presented work considers tissue-specific T_1_-corrections for each voxel (Wright et al., 2022), which showed a contrast shift for Glu and mI which are reported to have a strong T_1_-relaxation time tissue dependence (Wright, Murali-Manohar, et al., 2021). Within this work, statistical tests affirm that mI concentrations are increased in the occipital lobe WM compared to frontal lobe GM, occipital lobe GM, and parietal lobe GM; additionally, the parietal lobe WM concentrations are higher than the occipital lobe GM concentrations.

The herein obtained results generally agree with previous studies both at UHF and otherwise. Inter-subject CVs in Table 4 show that it is possible to confidently (CV < 10%) measure NAA, tCr, Glu, mI, tCho, Gln, GABA and Tau in a variety of major brain regions. While NAAG has a CV greater than 10% in all regions, metabolite maps have routinely showed NAAG spatially located in the WM (Bogner et al., 2012; Hangel et al., 2021; Henning et al., 2009; Nassirpour et al., 2016; Wright et al., 2022; Ziegs et al., 2023); acquiring more data could help to increase the confidence in NAAG concentration estimations.

More specifically, metabolite concentrations estimated in this study are in rough agreement with those previously acquired with the same methodology at 9.4 T (Wright et al., 2022). A noticeable shift is present, with many of the concentrations being slightly higher in this work. One notable characteristic in the concentration distributions (Figure 5) is an uneven density of concentrations biased towards lower values for metabolites with low peak intensities including GABA, NAAG, Scyllo or Asp. Recent and, so far, unpublished investigations have shown that this bias is caused by the LCmodel spectral fitting algorithm.

### 4.3 Limitations

This study aimed to achieve whole-brain coverage including cerebrum and cerebellum for metabolite mapping using a ^1^H FID-MRSI sequence; however, data quality was insufficient in lower slices to report reliable results. Thus, only cerebrum coverage was considered and yielded metabolite maps and regional concentrations for 12 metabolites. In general, data below slice 48 of the MNI space images which corresponds to below MRSI slice 10 were more prone to distortions that degraded metabolite image quality. A potential solution for this problem would be using an RF coil with improved longitudinal coverage and metabolite SNR in lower slices and most importantly improved B_0_ shimming (Motyka et al., 2019).

For illustrative purposes, Figure 7 and Figure 8 show the same coronal (15) and sagittal (25) voxel position across 12 transversal positions corresponding to respective MRSI slices acquired in one volunteer. As can be seen spectra in slices 1 – 9 (top to bottom) is good, the spectral line width starts to get very wide in slice 10 and after slice 10 the metabolite information is lost. For most volunteers, the data quality dropped similarly between slice 9 and slice 11. Improvements in B_0_ shimming and increased SNR in the lower slices of the brain could help to improve measurement quality and provide an avenue to investigate lower brain regions when part of full brain studies (Motyka et al., 2019).

**Figure 6:**
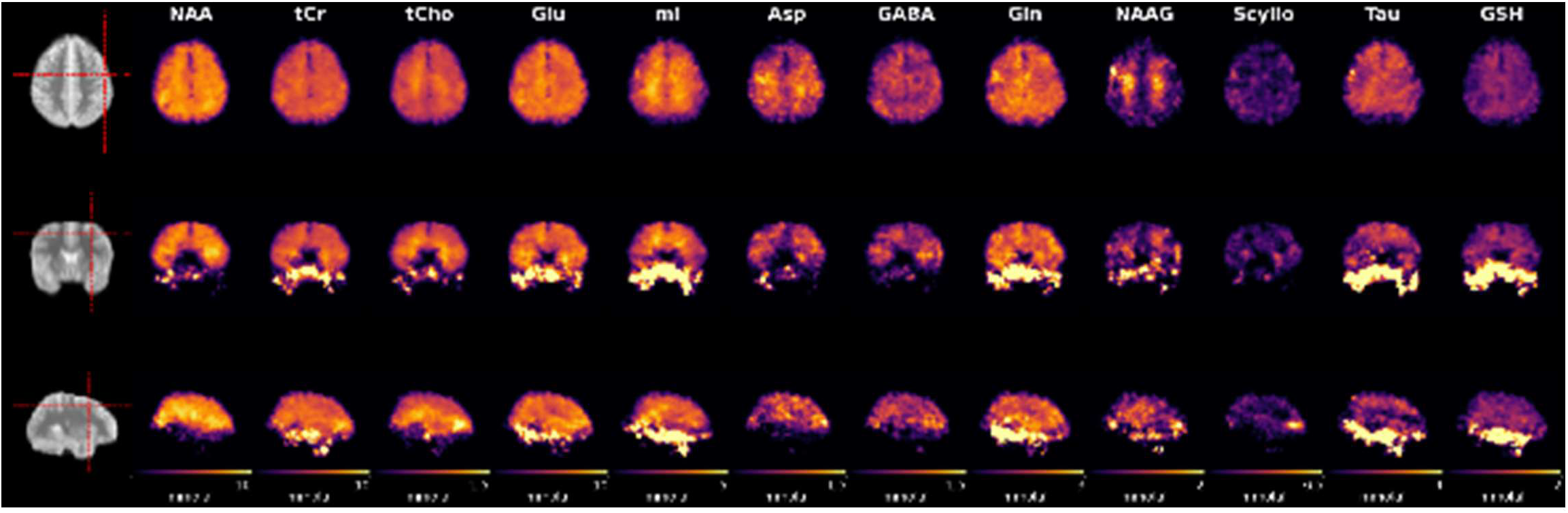
Transversal, coronal, and sagittal median metabolite maps [mmol kg^-1^]. Red dashed lines on the anatomical figures show the slices selected for display. All maps have 0 as the minimum and displayed with the max value for the maps noted on the color bars.

**Figure 7:**
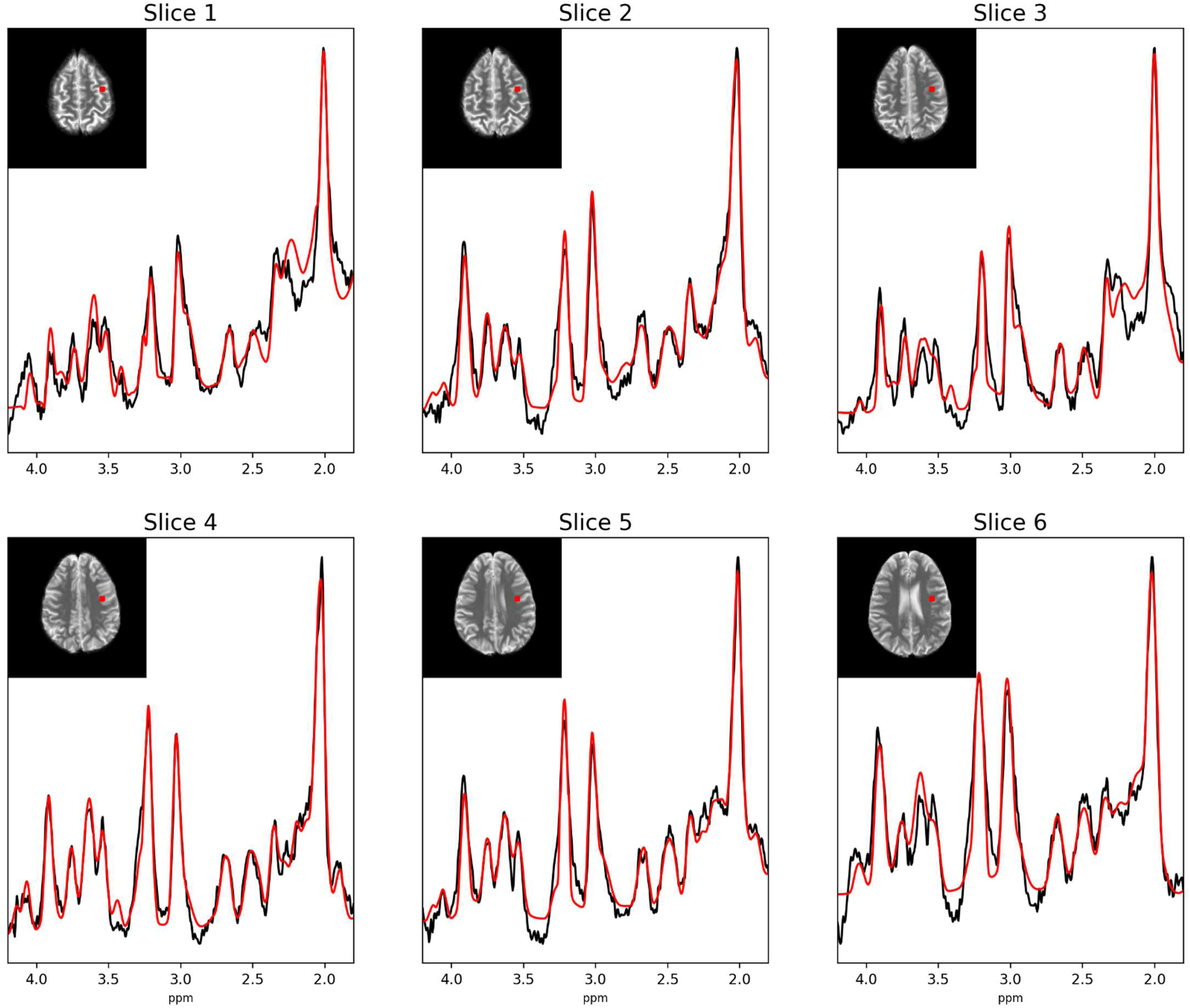
individual fitted voxels in the same matrix position (coronal: 15, sagittal: 24) across slices 1 to 6 from one volunteer. The red square shows the voxel position with roughly the same dimensions as the in vivo voxel.

**Figure 8:**
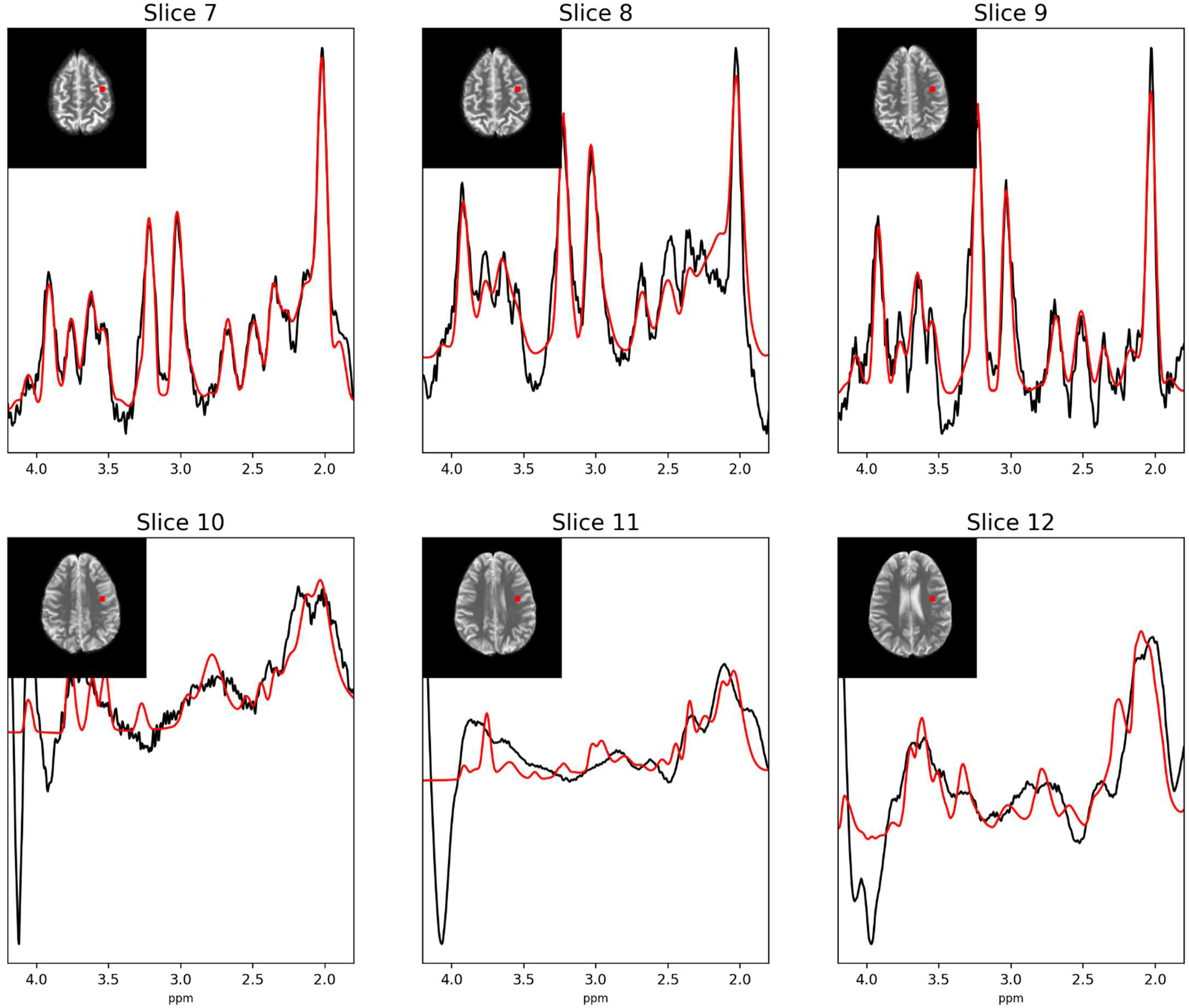
individual fitted voxels in the same matrix position (coronal: 15, sagittal: 24) across slices 7 to 12 from the same volunteer as in Figure 7. The red square shows the voxel position with roughly the same dimensions as the in vivo voxel. Note the dramatic shift in acquisition quality below slice 9 for this volunteer (not included in analysis).

In general, techniques such as motion correction (Moser et al., 2020) could dramatically reduce the CV for metabolite concentration estimates. Another strategy using MRSI could be to extract and sum region specific voxels in the step directly prior to fitting spectra (Schreiner et al., 2016, 2018). This could improve metabolite fitting performance in LCModel and therefore reduce inter-subject CVs.

From an analysis perspective, the WM atlas used to estimate WM concentrations is a tractography atlas, and the WM volumes do not include all WM tissue within brain regions. Developing a more appropriate atlas for MRSI studies could increase the number of WM voxels included in concentration estimates.

Finally, the inclusion of a larger number of volunteers in a future study could further increase the accuracy and precision of region specific metabolite concentration estimates in the entire brain.

## 5 Conclusion and outlook

The methods used in this study are a culmination of previous works that developed short TR, ^1^H FID MRSI at 9.4 T and reports group averaged quantitative ^1^H MRSI acquired in the human brain at 9.4 T with full cerebrum coverage for the first time. Co-registering MRSI data to the MNI space can positively contribute to further characterization of the concentrations and standardized reporting of the distributions of metabolites in the human brain. By reporting results in a standard space and applying full relaxation correction to the metabolite maps, future works would be able to characterize and compare regional spectra as well as combine metabolite maps across studies. Pooling ^1^H MRSI data from healthy volunteer and patient cohorts via this approach could bring more power to studies investigating differences between healthy and diseased populations in future and would also serve as a means to directly compare MRSI acquisition and quantification techniques.

