## Supporting Information for "Quantitative metabolic reference for healthy human cerebrum derived from group averaged 9.4T ^1^H MRSI data"

Supporting Information Table 1: combined anatomical regions from structural atlases to form major brain regions.

| **Frontal Lobe GM** |
| --- |
| • Frontal Pole |
| • Superior Frontal Gyrus |
| • Middle Frontal Gyrus |
| • Inferior Frontal Gyrus, pars triangularis |
| • Inferior Frontal Gyrus, pars opercularis |
| • Precentral Gyrus |
| • Frontal Medial Cortex |
| • Juxtapositional Lobule Cortex (formerly Supplementary Motor Cortex) |
| • Paracingulate Gyrus |
| • Cingulate Gyrus, anterior division |
| • Frontal Orbital Cortex |
| • Frontal Operculum Cortex |
| • Subcallosal Cortex |
| **Frontal Lobe WM** |
| • Anterior corona radiata |
| • Anterior limb of internal capsule |
| • Cingulum (cingulate gyrus) |
| • Superior fronto-occipital fasciculus (could be a part of anterior internal capsule) |
| • Uncinate fasciculus |
| **Insular Cortex** |
| • Insular Cortex |
| **Occipital Lobe GM** |
| • Lateral Occipital Cortex, superior division |
| • Lateral Occipital Cortex, inferior division |
| • Intracalcarine Cortex |
| • Cuneal Cortex |
| • Lingual Gyrus |
| • Occipital Fusiform Gyrus |
| • Supracalcarine Cortex |
| • Occipital Pole |
| **Occipital Lobe WM** |
| • Posterior corona radiata |
| • Posterior thalamic radiation (include optic radiation) |
| • Sagittal stratum (include inferior longitidinal fasciculus and inferior fronto-occipital fasciculus) |
| • Inferior fronto-occipital fasciculus |
| • Tapetum |
| **Parietal Lobe GM** |
| • Postcentral Gyrus |
| • Superior Parietal Lobule |
| • Supramarginal Gyrus, anterior division |
| • Supramarginal Gyrus, posterior division |
| • Angular Gyrus |
| • Cingulate Gyrus, posterior division |
| • Precuneous Cortex |

- Parietal Operculum Cortex

**Parietal Lobe WM**

- Superior longitudinal fasciculus

**Temporal Lobe**

- Temporal Pole
- Superior Temporal Gyrus, anterior division
- Superior Temporal Gyrus, posterior division
- Middle Temporal Gyrus, anterior division
- Middle Temporal Gyrus, posterior division
- Middle Temporal Gyrus, temporooccipital part
- Inferior Temporal Gyrus, anterior division
- Inferior Temporal Gyrus, posterior division
- Inferior Temporal Gyrus, temporooccipital part
- Parahippocampal Gyrus, anterior division
- Parahippocampal Gyrus, posterior division
- Temporal Fusiform Cortex, anterior division
- Temporal Fusiform Cortex, posterior division
- Temporal Occipital Fusiform Cortex
- Central Opercular Cortex
- Planum Polare
- Heschl's Gyrus (includes H1 and H2)
- Planum Temporale

**Corpus Callosum**

- Genu of corpus callosum
- Body of corpus callosum
- Splenium of corpus callosum

**Subcortical Structures**

- Thalamus
- Caudate
- Left Putamen
- Pallidum
- Hippocampus
- Amygdala
- Accumbens
- Brain-Stem

Supporting Information Figure 1: anatomical masks presented as the sum of the masks in sagittal, coronal, and transversal views. Note, the insular cortex and subcortical structures are not included in concentration estimates.

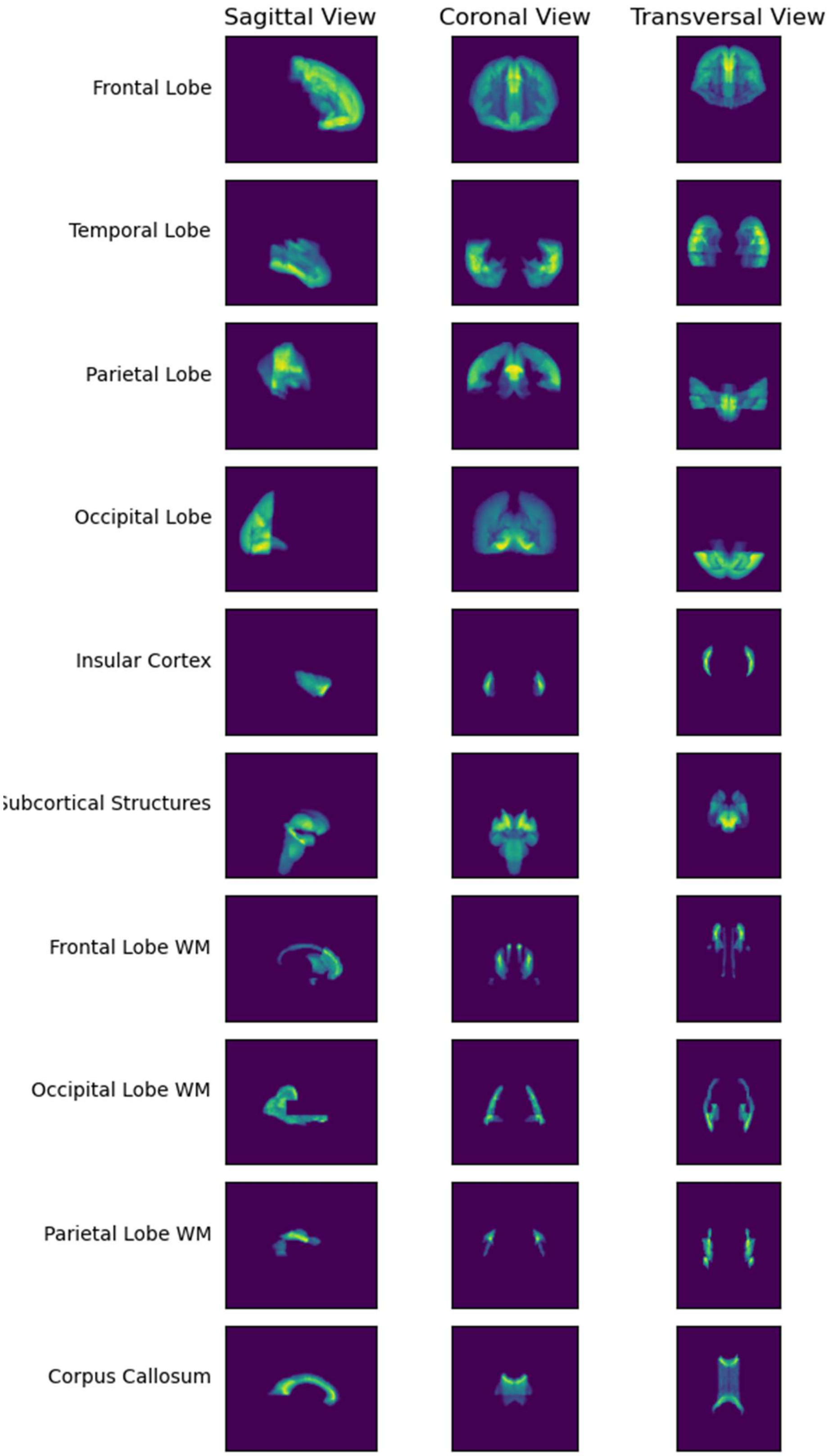

Supporting Information Figure 2: Median metabolite maps from eight volunteers in mMolar quantities.

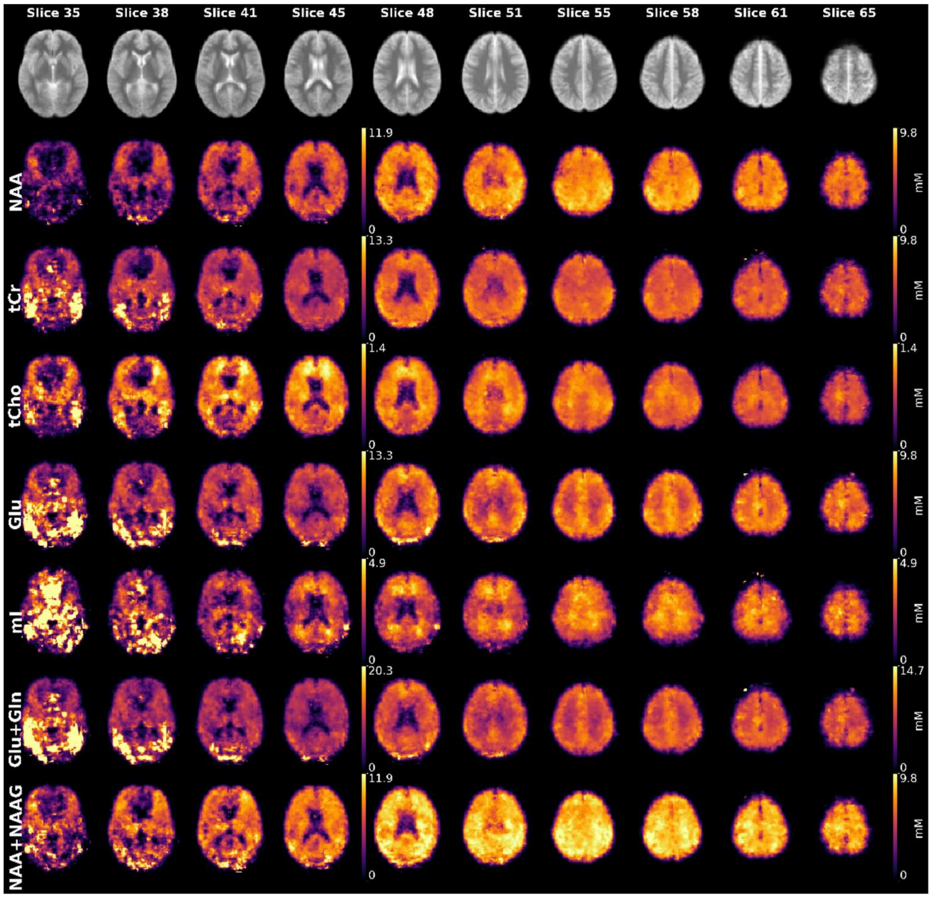

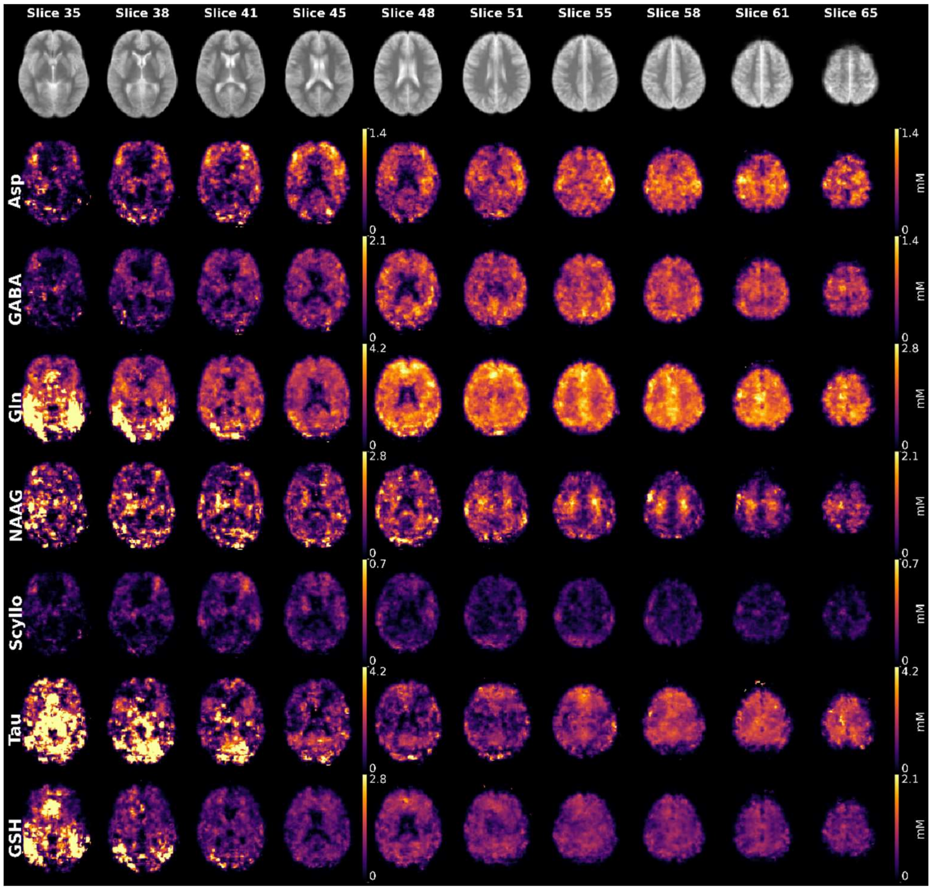

Supporting Information Figure 3: averaged CRLB maps with the CRLB threshold applied. These maps show how CRLB thresholds were applied to the metabolite maps prior to concentration estimates.

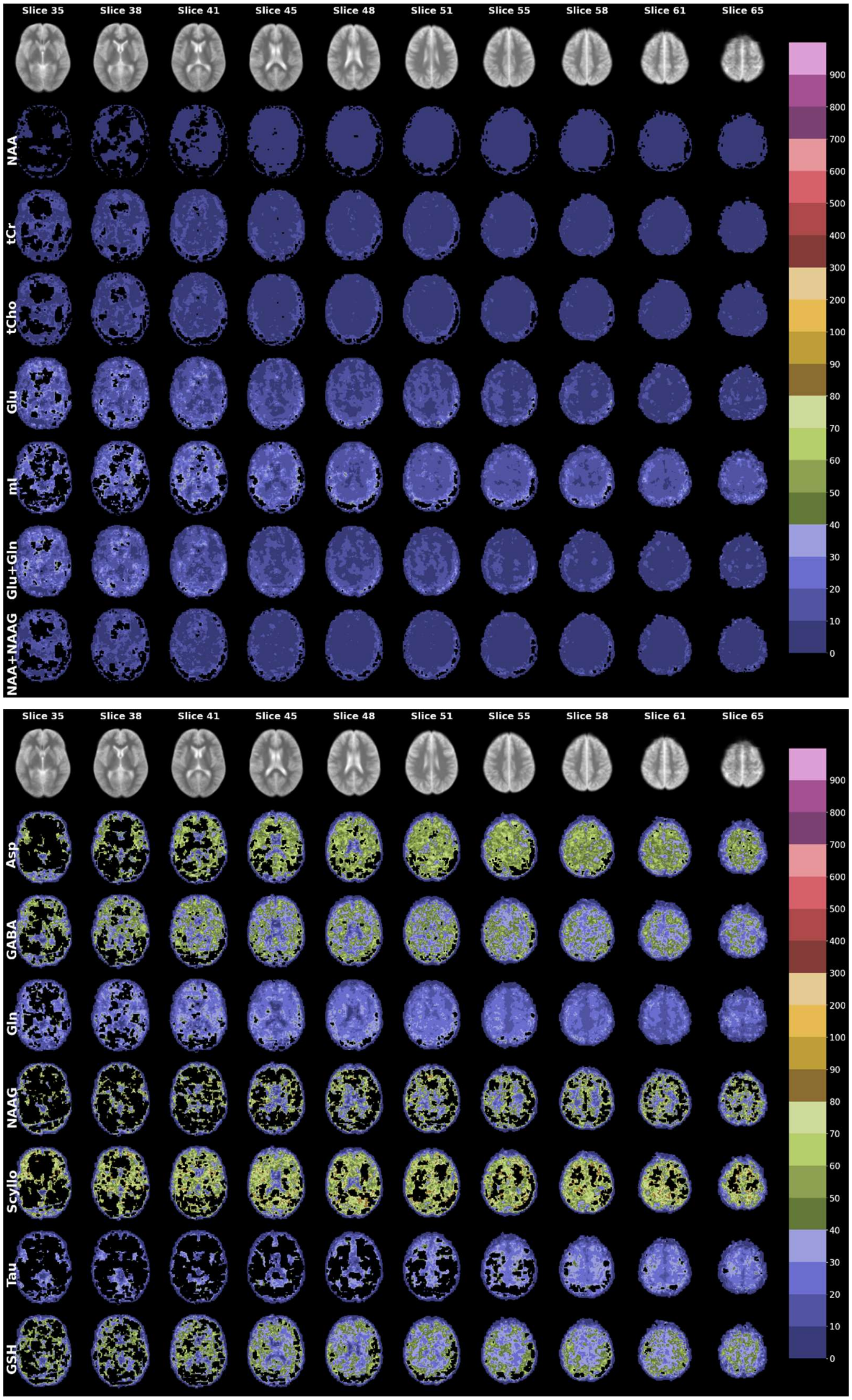

Supporting Information Table 2: Metabolite concentrations [mM]. Metabolite concentrations were calculated with the following:

$$[\mathrm{M}]_{\mathrm{molar}} = \frac{\mathrm{S_{met}} \times \left(\mathrm{f_{GM}} \times \mathrm{R_{H_2O_{GM}}} \times d_{GM} + \mathrm{f_{WM}} \times \mathrm{R_{H_2O_{WM}}} \times d_{WM} + \mathrm{f_{CSF}} \times \mathrm{R_{H_2O_{CSF}}} \times d_{CSF}\right)}{\mathrm{S_{H_2O}}(1 - \mathrm{f_{CSF}}) \times \mathrm{R_{met}}} \times \frac{\mathrm{N_{H_2O}}}{\mathrm{N_{met}}} \times [\mathrm{H_2O}]_{molal}$$

With the following variable definitions:

- S: observed signal
- f: fraction of a given tissue compartment
- R: relaxation state for a given tissue compartment
- d: visible water fraction of a given tissue compartment
- N: number of protons contributing
- $[H_20]_{molal}$: 55510 mol $kg^{-1}$

**Metabolite Concentrations [mM]**

| | Frontal Lobe GM | Temporal Lobe GM | Parietal Lobe GM | Occipital Lobe GM | Frontal Lobe WM | Occipital Lobe WM | Parietal Lobe WM | Corpus Callosum |
|---|---|---|---|---|---|---|---|---|
| **NAA** | 7.19 ± 1.32 | 7.7 ± 1.25 | 7.78 ± 1.17 | 7.93 ± 1.54 | 7.66 ± 0.91 | 7.5 ± 1.09 | 8.19 ± 0.82 | 6.32 ± 1.61 |
| **tCr** | 5.69 ± 1.7 | 6.66 ± 2.43 | 6.31 ± 1.31 | 6.03 ± 1.93 | 6.58 ± 1.33 | 5.89 ± 1.07 | 6.26 ± 0.85 | 4.95 ± 1.59 |
| **Glu** | 6.3 ± 2.02 | 7.61 ± 3.71 | 6.7 ± 1.72 | 6.37 ± 2.44 | 6.35 ± 1.59 | 5.4 ± 1.56 | 5.99 ± 1.47 | 4.94 ± 1.66 |
| **mI** | 3.0 ± 1.07 | 3.28 ± 1.99 | 3.01 ± 0.88 | 2.62 ± 0.89 | 3.47 ± 0.95 | 3.63 ± 1.08 | 3.17 ± 0.75 | 3.04 ± 1.11 |
| **tCho** | 0.81 ± 0.31 | 0.88 ± 0.33 | 0.85 ± 0.21 | 0.74 ± 0.32 | 1.13 ± 0.23 | 1.05 ± 0.19 | 1.06 ± 0.14 | 0.8 ± 0.34 |
| **Gln** | 1.73 ± 0.92 | 2.77 ± 2.69 | 1.85 ± 0.79 | 1.65 ± 1.09 | 2.07 ± 0.55 | 1.79 ± 0.73 | 1.83 ± 0.48 | 1.46 ± 0.77 |
| **NAAG** | 1.08 ± 0.79 | 2.18 ± 2.05 | 1.06 ± 0.66 | 1.45 ± 1.27 | 1.46 ± 0.67 | 1.43 ± 0.63 | 1.55 ± 0.54 | 1.26 ± 0.84 |
| **Asp** | 0.79 ± 0.54 | 0.69 ± 0.61 | 0.86 ± 0.5 | 0.7 ± 0.6 | 1.02 ± 0.27 | 0.82 ± 0.39 | 1.11 ± 0.28 | 0.58 ± 0.41 |
| **GABA** | 0.64 ± 0.4 | 0.65 ± 0.55 | 0.74 ± 0.4 | 0.66 ± 0.52 | 0.9 ± 0.33 | 0.71 ± 0.32 | 0.92 ± 0.3 | 0.55 ± 0.37 |
| **GSH** | 0.71 ± 0.41 | 1.87 ± 2.16 | 0.74 ± 0.35 | 0.68 ± 0.53 | 1.02 ± 0.33 | 0.79 ± 0.29 | 0.8 ± 0.21 | 0.65 ± 0.39 |
| **Tau** | 1.86 ± 0.99 | 11.06 ± 14.12 | 1.95 ± 0.79 | 1.94 ± 1.59 | 2.18 ± 0.43 | 1.84 ± 0.74 | 2.0 ± 0.37 | 1.42 ± 0.88 |
| **Scyllo** | 0.24 ± 0.1 | 0.29 ± 0.13 | 0.24 ± 0.1 | 0.29 ± 0.14 | 0.3 ± 0.12 | 0.24 ± 0.09 | 0.32 ± 0.12 | 0.22 ± 0.09 |

**Supporting Information Table 2:** Metabolite concentrations [mM] for the selected eight major brain regions. Regional masks can be found in Supporting Information Figure 1.

Supporting Information Table 3: Metabolite concentrations ratios (1/tCr). Metabolite concentrations were calculated with the following:

| Metabolite tCr Ratios | | | | | | | | |
|---|---|---|---|---|---|---|---|---|
| | **Frontal Lobe GM** | **Temporal Lobe GM** | **Parietal Lobe GM** | **Occipital Lobe GM** | **Frontal Lobe WM** | **Occipital Lobe WM** | **Parietal Lobe WM** | **Corpus Callosum** |
| **NAA/tCr** | 1.19 ± 0.15 | 1.07 ± 0.25 | 1.2 ± 0.13 | 1.19 ± 0.16 | 1.12 ± 0.15 | 0.99 ± 0.21 | 1.2 ± 0.16 | 1.27 ± 0.16 |
| **Glu/tCr** | 1.14 ± 0.2 | 1.18 ± 0.52 | 1.11 ± 0.16 | 1.14 ± 0.32 | 1.14 ± 0.22 | 1.04 ± 0.32 | 1.02 ± 0.22 | 0.94 ± 0.18 |
| **mI/tCr** | 0.51 ± 0.17 | 0.49 ± 0.48 | 0.49 ± 0.13 | 0.41 ± 0.14 | 0.44 ± 0.12 | 0.64 ± 0.46 | 0.56 ± 0.19 | 0.64 ± 0.16 |
| **tCho/tCr** | 0.15 ± 0.02 | 0.14 ± 0.03 | 0.14 ± 0.02 | 0.13 ± 0.02 | 0.15 ± 0.02 | 0.15 ± 0.03 | 0.18 ± 0.03 | 0.18 ± 0.02 |
| **Gln/tCr** | 0.35 ± 0.11 | 0.76 ± 0.62 | 0.34 ± 0.1 | 0.39 ± 0.19 | 0.37 ± 0.12 | 0.4 ± 0.22 | 0.33 ± 0.09 | 0.33 ± 0.11 |
| **NAAG/tCr** | 0.21 ± 0.17 | 0.28 ± 0.32 | 0.21 ± 0.13 | 0.34 ± 0.34 | 0.31 ± 0.18 | 0.37 ± 0.33 | 0.24 ± 0.11 | 0.27 ± 0.12 |
| **Asp/tCr** | 0.14 ± 0.08 | 0.08 ± 0.09 | 0.15 ± 0.07 | 0.13 ± 0.1 | 0.13 ± 0.06 | 0.1 ± 0.09 | 0.16 ± 0.04 | 0.16 ± 0.05 |
| **GABA/tCr** | 0.12 ± 0.06 | 0.09 ± 0.09 | 0.13 ± 0.05 | 0.12 ± 0.08 | 0.13 ± 0.06 | 0.12 ± 0.08 | 0.14 ± 0.04 | 0.13 ± 0.04 |
| **GSH/tCr** | 0.14 ± 0.05 | 0.49 ± 0.47 | 0.13 ± 0.04 | 0.16 ± 0.11 | 0.16 ± 0.06 | 0.28 ± 0.23 | 0.16 ± 0.04 | 0.15 ± 0.04 |
| **Tau/tCr** | 0.38 ± 0.16 | 1.35 ± 1.47 | 0.37 ± 0.1 | 0.6 ± 0.48 | 0.42 ± 0.14 | 1.65 ± 1.39 | 0.4 ± 0.08 | 0.41 ± 0.13 |
| **Scyllo/tCr** | 0.03 ± 0.02 | 0.02 ± 0.02 | 0.03 ± 0.02 | 0.03 ± 0.02 | 0.03 ± 0.02 | 0.03 ± 0.02 | 0.04 ± 0.02 | 0.03 ± 0.02 |

**Table 3:** Metabolite tCr ratios for the selected eight major brain regions. Regional masks can be found in Supporting Information Figure 1.

Supporting Information Table 4: Metabolites where the concentration difference yielded a corrected p-value < 0.05 are shown at the intersection of anatomical regions. This figure serves as a compliment to Figure 4 in the main text.

| **Regions where corrected p-values < 0.05** | | | | | | | | |
|---|---|---|---|---|---|---|---|---|
| **Region** | **Metabolites** | | | | | | | |
| *Corpus Callosum* | | | | | | | | |
| *Frontal Lobe GM* | | | | | | | | |
| *Frontal Lobe WM* | NAA, tCr, GABA, GSH, Scyllo, Glu+Gln | NAA, tCr, tCho, Asp, GABA, GSH, Scyllo, NAA+NAAG | | | | | | |
| *Occipital Lobe GM* | | | tCho, Asp, GABA | | | | | |
| *Occipital Lobe WM* | NAA | NAA, tCr, tCho, Glu, mI, NAA+NAAG | | tCho, mI | | | | |
| *Parietal Lobe GM* | | tCr | tCho, GABA, GSH, Scyllo | | tCho, Glu, mI | | | |
| *Parietal Lobe WM* | NAA, GABA, Scyllo, NAA+NAAG | NAA, tCr, tCho, GABA, Scyllo, NAA+NAAG | | NAA, tCho, Asp, GABA, mI, NAA+NAAG | | NAA, tCho, Scyllo, NAA+NAAG | | |
| *Temporal Lobe GM* | Glu, Glu+Gln | tCr, GSH, Glu+Gln | NAA, tCho, Asp, GABA | | tCho, Glu, Glu+Gln | | NAA, Asp, GABA, NAA+NAAG | |
| **Region** | *Corpus Callosum* | *Frontal Lobe GM* | *Frontal Lobe WM* | *Occipital Lobe GM* | *Occipital Lobe WM* | *Parietal Lobe GM* | *Parietal Lobe WM* | *Temporal Lobe GM* |

MRSinMRS methods table

| Hardware | |
|---|---|
| Field strength | 9.4 T |
| Manufacturer | Siemens |
| Model | Magnetom |
| RF Coil | 18Tx/32Rx phased array coil (Avdievich et al., 2018) |
| **Acquisition** | |
| Pulse sequence | FID-MRSI |
| Repetition time, TR | 300 ms |
| Acquisition delay, TE* | 1.5 ms |
| Flip angle | 47° |
| Slice coverage | cerebrum |
| MRSI grid size | Variable to achieve more slice coverage within a shorter time frame. |
| Nominal voxel size | 6 x 6 x 6 $mm^3$ |
| Water suppression method | Three unmodulated Hanning-filtered Gaussian pulses (BW = 180 Hz, duration = 5 ms) with flip angles of 90°, 79.5°, and 159°; the inter-pulse delay between all pulses was 20 ms. |
| Shimming method | 2nd-order vendor implemented shimming |
| **Analysis method** | |
| Fitting software | LCModel v6.3-1L |
| Quantification | Voxel-specific $T_1$-corrections |
| Coregistration to MNI152 | FSL FLIRT |
| Brain region masking | 1. Harvard-Oxford maximum probability cortical atlas [2 mm]<br>2. Harvard-Oxford maximum probability subcortical atlas [2 mm]<br>3. Johns Hopkins University, International Consortium for Brain Mapping template [2mm] |
